# Unzipped assemblies of polyploid root-knot nematode genomes reveal new kinds of unilateral composite telomeric repeats

**DOI:** 10.1101/2023.03.29.534350

**Authors:** Ana-Paula Zotta Mota, Georgios D Koutsovoulos, Laetitia Perfus-Barbeoch, Evelin Despot-Slade, Karine Labadie, Jean-Marc Aury, Karine Robbe-Sermesant, Marc Bailly-Bechet, Caroline Belser, Arthur Péré, Corinne Rancurel, Djampa K Kozlowski, Rahim Hassanaly-Goulamhoussen, Martine Da Rocha, Benjamin Noël, Nevenka Mestrovic-Radan, Patrick Wincker, Etienne GJ Danchin

## Abstract

Telomeres play central roles in senescence, aging and chromosome integrity. Using ONT long read sequencing we have assembled the genomes of *Meloidogyne incognita*, *M. javanica* and *M. arenaria*, the three most devastating plant-parasitic nematodes at unparalleled contiguity. The telomeric repeat (TTAGGC)n, evolutionarily conserved in nematodes, was not found in these genomes. Furthermore, no evidence for a telomerase enzyme or for orthologs of *C. elegans* telomere-associated proteins could be found. Instead, we identified species-specific composite repeats mostly present at one end of contigs. These repeats were G-rich, oriented and transcribed, similarly to known telomeric repeats. Using FISH we confirmed these repeats were present at one single end of *M. incognita* chromosomes. The discovery of a new kind of telomeric repeat in these species highlights the evolutionary diversity of chromosome protection systems despite their central roles and opens new perspectives towards the development of more specific control methods against these pests.

## Introduction

Telomeres are nucleoprotein complexes that cap and protect eukaryotic linear chromosomes. They play multiple central cellular functions such as protection against chromosomal fusions or avoidance of recognition of chromosome ends by DNA damage response pathways. They also ensure no loss of genetic information occurs during DNA replication and are involved in aging and senescence. Their misregulation or malfunction can lead to severe disease including uncontrolled ‘immortal’ cell proliferation and cancers (Revy *et al*., 2023).

Telomeres possess an ensemble of evolutionarily conserved features considered as canonical signatures. First, telomeric DNA is usually constituted by terminal short and simple G-rich repeats at chromosome ends and can fold in G4-quadruplex secondary structures. Second, to compensate for their shortening at each replication, these repeats are usually re-multiplied at chromosome ends by a telomerase enzyme. Telomerase is a RNA-dependent polymerase composed of a telomerase RNA component (the repeat template) and a catalytic subunit, telomerase reverse transcriptase which copies this template. Third, telomeric DNA usually interacts with a complex of proteins (shelterin) and telomeric RNA (TERRA) that play diverse important roles in their stability and function.

Although this system is considered canonical in eukaryotes, including in human beings, some exceptions exist (Louis, 2002). For instance, *Diptera* insects seem to have ancestrally lost the telomerase enzyme (Mason *et al*., 2016). Consistent with the absence of telomerase, simple telomeric repeats are not found at chromosome ends in several *Diptera* genomes. Some species display telomeric retrotransposons either instead of simple telomeric repeats or in association with them (Kordyukova *et al*., 2018; Louis, 2002). For example, in *Drosophila melanogaster*, three different retrotransposons, distantly related to the Jockey superfamily and specifically copied and pasted at chromosome ends by reverse transcription, play the role of telomeric DNA (Casacuberta, 2017; Pardue and DeBaryshe, 2013). Besides, other *Diptera*, such as *Chironomus thummi* mosquitoes, display complex repeats with no evident similarity to retrotransposons and their mechanism of multiplication remains unsolved. (Martinez *et al*., 2001).

In nematodes, it has been assumed that a (TTAGGC)n DNA repeat multiplied by a telomerase is the canonical system. Indeed, the repeat and the enzyme are both evolutionarily conserved between distant nematode species, including in *C. elegans* (Carlton *et al*., 2022). The telomerase enzyme is encoded by the protein-coding gene trt-1 in *C. elegans* but the RNA template of the repeat has not been identified so far (Carlton *et al*., 2022). In the animal parasite *Parascaris univalens*, a non G-rich motif (TTGCA)n was initially described to be repeated albeit degenerated and to play the role of telomeric DNA in the somatic genome of this species (Teschke *et al*., 1991). However, a more recent study showed that the germline chromosomes of this species are in fact capped by terminal (TTAGGC)n repeats that follow the (TTGCA)n and other more degenerate repeats (Niedermaier and Moritz, 2000). Selective DNA elimination removes all germline telomeric repeats and other satellite DNA in somatic cells. Fluorescent in-situ hybridization showed that (TTAGGC)n telomere repeats are afterward re-generated at the extremities of all the somatic chromosomes. Analysis of the genome of the uni-chromosomal nematode *Diploscapter pachys* revealed no evidence for the canonical nematode telomeric repeats, even in the raw genome reads (Fradin *et al*., 2017). The same paper suggested that the canonical nematode repeat was also absent from the genomes of the vertebrate parasite *Trichinella spiralis* as well as the root-knot nematode *Meloidogyne hapla*. These observations suggest that different repeats could protect chromosome ends in these species and the (TTAGGC)n repeat might not be as universal as assumed in nematodes.

Shelterin proteins form bridged telomeric protein complexes and are involved in many telomeric functions like telomerase regulation, protection against activation of the DNA damage response (DDR), chromosome stability, long and short range transcriptional regulation and meiosis (Giraud-Panis *et al*. 2018). The shelterin complex is usually composed of both single-stand and double-strand telomeric DNA binding proteins as well as other proteins that bridges these proteins. In humans and ciliates, there is only one POT1, a single-stranded telomere binding protein (Protection of Telomeres 1) with several oligosaccharide/oligonucleotide binding folds (OB folds). OB folds found across distant organisms commonly bind single stranded DNA, although a wider specificity, including to other ligands was shown in some cases. The conserved OB-fold of POT1 is necessary for its single stranded DNA binding specificity and the telomere protective functions. In *C. elegans*, there are 4 distinct POT1 homologs containing only one OB fold: POT-1, POT-2, POT3 and MRT-1 (Carlton *et al*., 2022). POT1 and POT-2 exhibit structural similarity to the first and second OB fold respectively of the mammalian telomere binding protein hPOT1. POT1 and POT-2 can promote T-loop formation in vitro (Raices *et al*., 2008) and repress telomerase activity in vivo in *C. elegans* (Shtessel *et al*., 2013). POT-3 exhibits a structural similarity to the second OB fold and was recently found to preferentially bind the terminal DNA-repeat on the telomeric G-rich 3’ overhang (Yu *et al*., 2023), while its absence leads to uncontrolled telomere lengthening. MRT-1, the fourth POT-1 homolog, possesses a SNM1 nuclease domain in addition to a N-teminal domain similar to the second OB-fold of POT1, and this gene is required for telomerase activity (Meier *et al*., 2009). The number of POT1 homologs in nematodes seems to vary (Carlton *et al*., 2022). TRF (Telomeric Repeat Factors) proteins and 3′-overhang-binding heterodimers are linked by a protein bridge, formed by Rap1 and Poz1 in the yeast *Schizosaccharomyces pombe* and by TIN2 (an ortholog of Poz1) in mammals. However, in *C. elegans*, TRF protein homologs with a TRFH (Telomeric Repeat Factors Homology) domain or a telobox at their C-terminus could not be found. Thus, the shelterin complex seems to be composed of other types of proteins. Two proteins in *C. elegans*, TEBP1/DTN-1 and TEBP2/DTN-2, were shown to bind double-stranded telomeric DNA and POT1 (Yamamoto *et al*., 2021). Sterility is observed after one or several generations in tebp-1; tebp-2 double mutants (Dietz *et al*., 2021; Yamamoto *et al*., 2021). A single TEBP homolog was found in many nematodes including in *C. briggsae* (Dietz *et al*., 2021).

Although the presence of a telomerase enzyme, a simple (TTAGGC)n telomeric repeat and a shelterin complex is considered the default system in nematodes, no extensive search has been performed in the phylum nematoda. Furthermore, because many nematode genomes are still highly fragmented and short-read sequencing technologies hardly resolve long repetitive regions, determination of DNA repeats at chromosome ends has remained difficult. Long-read sequencing technology, elected 2022 method of the year (Marx, 2023) opened new perspectives towards resolving the genomes of non-model species.

Here, using ONT long-read sequencing, we have assembled the genomes of the root-knot nematodes *Meloidogyne incognita*, *M. javanica* and *M. arenaria*, three major agricultural pests of major concern worldwide. Because these genomes have a complex polyploid structure, previous short-read versions of the assemblies of the same species remained quite fragmentary (Blanc-Mathieu *et al*., 2017; Szitenberg *et al*., 2017). Our long-read assemblies were two orders of magnitude more contiguous and allowed unzipping and assembling the three (*M. incognita*) and four (*M. javanica*, *M. arenaria*) subgenomes. We used these more accurate representations of the genomes to investigate how chromosomes start and end and whether the telomere system considered to be canonical in nematodes was conserved. We discovered new kinds of complex repeats enriched at contigs ends and so far specific to these species. Using FISH, we confirmed telomeric localization of these repeats in *M. incognita*. We also investigated how telomeric repeats, and proteins involved in telomere maintenance and functions were conserved across the nematoda phylum including in root knot nematodes revealing their absence in these species.

## Results

### Long-read sequencing improves genome assemblies and confirms polyploidy

We assembled the genomes of *M. incognita* (*Minc*), *M. javanica* (*Mjav*) and *M. arenaria* (*Mare*) in 291, 364 and 377 contigs, respectively, representing a massive improvement compared to previous versions of these genomes (Blanc-Mathieu *et al*., 2017), assembled in ca. 12,000 - 31,000 scaffolds (Table 1). The new assembly sizes reached 199.4, 297.8 and 304.3 Mb for *Minc*, *Mjav* and *Mare*, consistent with previous flow cytometry estimates, and the GC content was around 30% for all species (Table 1). For all three species, the contig N50 lengths were around 2Mb which is two orders of magnitude higher than the previous N50 values (0.01 - 0.04 Mb) and the highest for a publicly available *Meloidogyne* genome so far. Analysis of k-mer distribution in the sequencing reads (Supplementary Fig. 1) suggested a triploid (AAB) genome for Minc and tetraploid (AABB) genomes for Mjav and Mare, with respectively 6.6, 8.7 and 9.2% average nucleotide divergence between the homoeologous genome copies (Supplementary Fig. 2). These results were also consistent with previous analyses in the same species (Blanc-Mathieu *et al*., 2017; Jaron *et al*., 2021). Assessment of completeness via CEGMA (Parra *et al*., 2007) and BUSCO (Manni *et al*., 2021) genes shows a moderate improvement and suggests previous assemblies, albeit much more fragmented, were almost as complete as the new assemblies in terms of gene content (Table 1). Consistent with k-mer estimation of ploidy, CEGMA genes were present on average in 2.7 copies in Minc and 3.8 copies in Mjav and Mare. Supporting genome completeness, almost all k-mers present in the reads were also found in the assembly for the three species (Supplementary Fig. 3). Finally, analysis of contigs coverage, GC content and taxonomic assignment, revealed no evidence for contamination and allowed isolating contigs corresponding to the mitochondrial genome in each species (Supplementary Fig. 4).

**Table 1:** genome assembly metrics (a: previous versions from (Blanc-Mathieu *et al*., 2017) b: this study). CC: % of complete CEGMA genes, A: average number of orthologs per CEGMA gene. BC: % of complete BUSCO genes, D: % of duplicated complete BUSCO genes, F: % of fragmented BUSCO genes, M: % of missing BUSCO genes.

|  | *Minc*<sup>a</sup> | *Minc*<sup>b</sup> | *Mjav*<sup>a</sup> | *Mjav*<sup>b</sup> | *Mare*<sup>a</sup> | *Mare*<sup>b</sup> |
| --- | --- | --- | --- | --- | --- | --- |
| Flow cytometry<sup>a</sup> | 189 +/- 15 |  | 297 +/- 27 |  | 304 +/- 9 |  |
| Assembly size (Mb) | 183.5 | 199.4 | 235.8 | 297.8 | 258.1 | 304.3 |
| N50 (Mb) | 0.04 | 1.86 | 0.01 | 2.07 | 0.02 | 2.04 |
| # Contigs | 12,091 | 291 | 31,341 | 364 | 26,196 | 377 |
| CEGMA (N: 248) | CC: 94.76<br>A: 2.93 | CC: 94.76<br>A: 2.69 | CC: 92.74<br>A: 3.68 | CC: 95.56<br>A: 3.75 | CC: 93.95<br>A: 3.57 | CC: 95.97<br>A: 3.78 |
| BUSCO metazoa (N: 954) | BC:56.3<br>(D:45.0)<br>F:9.3 M:34.4 | BC:56.6<br>(D:47.6) F:9.1<br>M:34.3 | BC:56.8<br>(D:45.0)<br>F:8.9 M:34.3 | BC:57.1<br>(D:51.5)<br>F:8.9 M:34.0 | BC:56.6<br>(D:47.1) F:9.8<br>M:33.7 | BC:57.9<br>(D:51.8) F:8.6<br>M:33.5 |
| GC% | 29.75 | 30.11 | 29.96 | 30.20 | 29.97 | 30.29 |

Previous observations (Blanc-Mathieu *et al*., 2017; Jaron *et al*., 2021) as well as k-mer distribution and CEGMA results suggest a triploid genome for Minc and tetraploid genomes for Mjav and Mare. To further investigate the genome structures, we predicted genes with EuGene (Sallet *et al*., 2019) in these three genomes (Supplementary Table 1) and used them as anchors to detect duplications with McScanX (Wang *et al*., 2012).

The McScanX analysis (Table 2) revealed only 13.6, 9.3 and 8.9% of predicted protein-coding genes were singleton in the Minc, Mjav and Mare genomes, implying that the vast majority of the genes were duplicated (86 - 91%). Similar analyses on previous genome assemblies, also concluded the vast majority of protein-coding genes were duplicated but only allowed assigning 28.5, 4.6 and 3.4 % of the genes to the WGD category, in Minc, Mjav and Mare, respectively (Blanc-Mathieu *et al*., 2017). It was hypothesized the limitation was due to the high fragmentation and low N50 of previous assemblies. In the new, more contiguous assemblies, the majority of protein coding genes (66 - 75%) were classified in the category ‘whole genome duplication (WGD) / segmental duplication’, consistent with the previous hypothesis that the three genomes are polyploid. Within this category of duplications, we observed a peak of genes in blocks duplicated at a depth of 3X for Minc (54% of WGD) while for *Mjav* and *Mare* we observed a peak at a depth of 4X with respectively 65% and 69% of the genes present in the WGD category. These results are consistent with Minc being triploid while Mjav and Mare being tetraploid. For all the species, more than 60% of the contigs form duplicated blocks with other contigs. The rest of the genes were distributed in the categories ‘dispersed duplications’ (12.6 -15.1%), ‘proximal duplications’, meaning duplicated genes are separated by one to 5 other genes in the same contig (2.0 - 3.3%) and ‘tandem duplications’, corresponding to consecutive cis-duplications in the same contig (1.3 - 1.8%). Overall, the new assemblies based on long reads produced much more contiguous genomes with N50 values two orders of magnitude higher, allowing better resolution of the duplicated genome structure. With half of the contigs reaching several megabases for all the species, and the biggest probably nearly representing whole chromosomes, new questions such as how chromosomes start and end in these nematodes can now be investigated.

**Table 2:** classification of duplicated genes in the three Meloidogyne genomes.

| Feature / species | *Minc* | *Mjav* | *Mare* |
| --- | --- | --- | --- |
| Protein-coding genes | 52,182 | 61,060 | 59,500 |
| Singleton | 7,104 (13.6%) | 5,676 (9.3%) | 5,294 (8.9%) |
| Dispersed dup. | 7,854 (15.1%) | 7,760 (12.7%) | 7,507 (12.6%) |
| Proximal dup. | 1,711 (3.3%) | 1,277 (2.09%) | 1,195 (2.0%) |
| Tandem dup. | 957 (1.8%) | 767 (1.26%) | 836 (1.41%) |
| WGD / Segmental dup.<br>[Dup. depth peak] | 34,556 (66.2%)<br>[3X - 54%] | 45,580 (74.65%)<br>[4X - 65%] | 44,668 (75.1%)<br>[4X - 69%] |
| Aligned blocks | 730 | 1,041 | 891 |
| Contigs involved | 193 (66.3%) | 246 (67.6%) | 232 (61.5%) |

### *Meloidogyne* genomes lack canonical nematode telomere repeats and most telomere-associated proteins known in *C. elegans*

#### Telomeric repeats

The chromosomes of *C. elegans* and most nematodes start and end with an evolutionarily conserved (TTAGGC)n telomeric repeat (Carlton *et al*., 2022). In the *Minc, Mjav*, and *Mare* genome assemblies generated in our study, this nematode telomeric sequence was found repeated maximum two times consecutively, and never at contig ends. Furthermore, this telomeric sequence was also not found repeated at contig or scaffold ends in any of the publicly available *Meloidogyne* genome assemblies. To better investigate how evolutionarily conserved and thus how canonical the *C. elegans* telomeric repeat is in the phylum nematoda, we searched in all the available nematode genomes assembled with a minimal N50 length of 100kb. In contrast to the *Meloidogyne* genus, (TTAGGC)n could be identified repeated in arrays in the genome assemblies for most other nematode genera investigated, including in *Pratylenchus penetrans*, the most closely related species with a genome available (Figure 1, Supplementary Table 2). In the rest of nematode clade IV, to which the *Meloidogyne* belong, the canonical telomeric repeat was well conserved except in the *Strongyloididae* group of animal parasites. Overall, ancestral state reconstruction using parsimony, suggests the *C. elegans* telomere repeat was present in the last common ancestor of nematodes but lost in *Trichinellidae* (clade I), in *Diploscapter* (clade V) and among Clade IV in the *Strongyloididae* and *Meloidogyne* groups (Supplementary Fig. 5). Further search for any arrays of perfectly conserved nucleotide repeats of size 6 to 12 at contig ends of *Minc*, *Mjav* and *Mare* also failed to identify any simple telomere repeats in these species.

**Figure 1:**
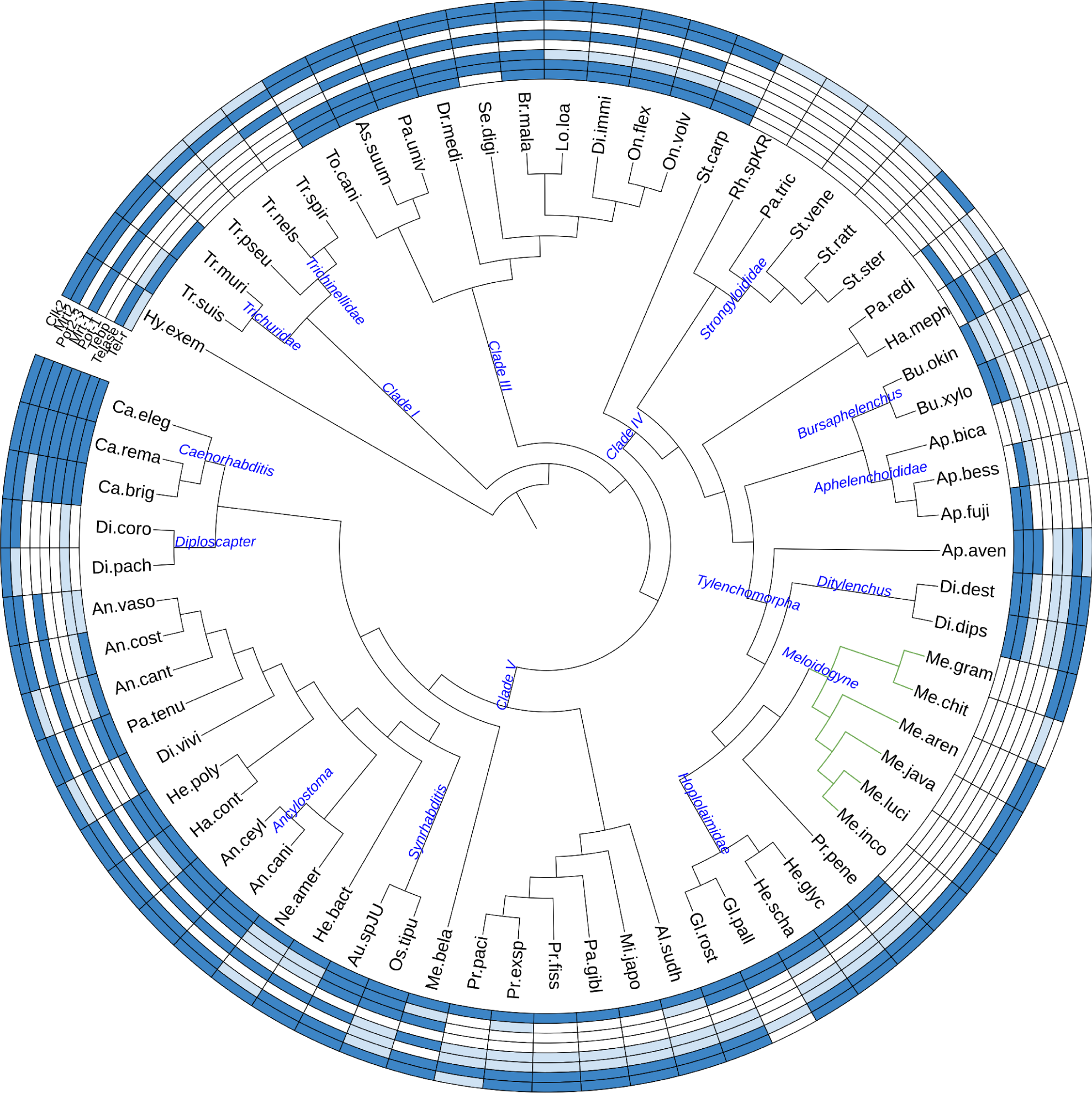
distribution of telomere repeats and telomere-associated proteins in the phylum nematoda. Blue color means present, light-blue means partial evidence and blank no evidence. Tel-r: *C. elegans* telomeric repeat (TTAGGC)n except for *Hypsibus exemplaris* in which another simple 9-nucleotides repeat has been described (Yoshida *et al*., 2017). Telase: telomerase reverse transcriptase. Tebp: ds-telomeric DNA binding proteins Tebp1 or Tebp2. Pot1: ss-telomeric DNA binding protein Pot-1. Mrt1: ss-telomeric DNA binding protein Mrt-1. Pot2-3: ss-telomeric DNA binding proteins Pot2 or Pot3. Mrt2 and Clk2: proteins with a putative function in telomere length regulation. In green lines, *Meloidogyne* genus.

#### Telomerases

Because no evidence for the canonical *C. elegans* telomeric repeats or other simple 6-12mer repeat arrays were found at contig ends in the *Meloidogyne* genomes, we investigated whether a telomerase enzyme was present in these and other nematode genomes. Combined search of protein motifs, orthology relationship at the protein level and homology search in the nematode genomes revealed no evidence for a telomerase enzyme in any of the *Meloidogyne* genomes investigated, not even at the pseudogene level and this absence included the three genomes sequenced in this study. This observation is consistent with the absence of the canonical nematoda telomeric repeat (TTAGGC)n or any other simple tandem repeats at contig ends in these species. Within clade IV, besides the *Meloidogyne*, evidence for telomerase enzymes were found in most other nematodes, including in *Pratylenchus penetrans*, the closest outgroup species relative to the *Meloidogyne* with available genome data and most other *Tylenchomorpha* (Figure 1, Supplementary Table 2). However, no evidence for a telomerase enzyme could be found in the *Strongyloididae*, which also showed no evidence for the *C. elegans* telomere repeat. Extending the analysis to all the other nematode clades indicated that telomerase enzymes were found in all species in clade III. However, in clade V, although the *Caenorhabditis*, *Diploscapter* and *Ancylostoma* genera all had evidence for a telomerase, the rest of genera either displayed sparse evidence or no evidence for a telomerase. Finally, in clade I, no species presented convincing evidence supporting the presence of a telomerase enzyme. Ancestral state reconstruction suggested a telomerase enzyme was present in the nematode ancestor but secondarily lost multiple times in the *Meloidogyne*, in *Strongyloididae*, in the *Trichinellidae* and *Diplogastridae* (Supplementary Fig. 6). The absence of detectable telomerase suggests the presence of an alternative lengthening of telomere (ALT) pathway in *Meloidogyne*.

#### Shelterin proteins

No evidence for proteins or even a domain interacting with single-stranded telomeric repeats could be identified in any *Meloidogyne* species (Figure 1, Supplementary Table 2). In the rest of clade IV nematodes, some species had evidence for a ssDNA-binding domain of telomere protection protein, including *Pratylenchus penetrans* as well as other groups of plant-parasitic nematodes such as *Hoplolaimidae* (*Globodera* and *Heterodera*), *Ditylenchus* or *Bursaphelenchus*. However, these proteins had no recognizable orthology relationships with those of *C. elegans*. This suggests orthologs of *C. elegans* proteins have highly diverged or other proteins might have been recruited to play the same role in these nematodes. In clade I and clade III nematodes, single-stranded telomeric DNA-binding domains were found but orthology evidence could be identified only for Mrt-1, suggesting the rest of POT-1 homologs were absent. Finally, in clade V, the four POT-1 homologs could be identified in *C. elegans* and *C. remanei* but only two of them (Mrt-1 and Pot-1) in *C. briggsae*. In the rest of nematodes in this clade, evidence for a ssDNA-binding domain of telomere protection proteins could be identified in a majority of species. Yet, orthology relationships could be confirmed only for Mrt-1 in a few species and several species showed no evidence for the presence of any of the four POT-1 homologs (e.g. *Diploscapter* genus). This ensemble of observations supports the idea that the number of POT-1 homologs varies in nematodes.

Similarly to single-strand telomere-binding proteins (i.e. POT-1 homologs), no evidence for homologs of *C. elegans* double-strand telomeric DNA-binding proteins or of presence of the corresponding domain could be found in any of the *Meloidogyne* genomes (Figure 1, Supplementary Table 2). Similarly, no evidence for the presence of these proteins or a dsDNA-binding domain of telomere-associated protein was identified in other nematodes from clade IV, to the exception of *Aphelenchus avenae* and *Halicephalobus mephisto*. No nematode species from clade I showed evidence for the presence of tebp-1 or tebp-2 protein domains or homologs. In contrast, a majority of nematode species from clade III showed evidence for homologs of these proteins. Finally, in clade V, apart from the *Caenorhabditis*, the *Synrhabditis* and *Ancylostoma* groups, in which tebp homologs could be identified, most other genera also showed either no or sparse evidence for conservation of these proteins or associated domain.

#### Other proteins related to telomere maintenance or elongation

Besides single and double stranded telomeric DNA-binding proteins which can be considered equivalent to the shelterin complex in *C. elegans*, other proteins are described as playing a role in telomere maintenance or elongation.

MRT-2 is a homolog of yeast rad17 and human rad1 DNA damage checkpoint proteins, and *C. elegans* mutants of this gene show defect in germline immortality associated with telomere shortening (Ahmed and Hodgkin, 2000). It has been proposed that MRT-2 might regulate telomere length but not necessarily directly bind telomeric DNA (Dietz *et al*., 2021). Our study shows MRT-2 is widely conserved in nematodes, including in the *Meloidogyne* species, as well as all clade I and clade III species. This is supported both by the presence of the protein domain and orthology relationship (Figure 1, Supplementary Table 2). Within clade IV, MRT-2 protein domains seem to be absent from the plant-parasitic *Aphelenchoididae* as well as from *Strongyloididae* animal parasites. In clade V, MRT-2 was conserved in 20 of the 25 species, covering all nematode superfamilies in this clade. Overall, the strong conservation of MRT-2, even in species that display neither canonical telomeres nor telomerase enzymes, suggests that besides the putative role in telomere length regulation, this protein plays a core central role in nematode DNA damage checkpoint.

CLK-2 is an ortholog of human TELO-2 telomere maintenance protein, which is essential for viability across eukaryotic evolution from yeast to vertebrate (Moser *et al*., 2009). Although a role in telomere length regulation has initially been proposed (Bénard *et al*., 2001; Lim *et al*., 2001), the exact function of clk-2 relative to telomeres remains unclear because mutants display pleiotropic effects and it was also suggested that clk-2 is mainly a DNA damage checkpoint protein (Ahmed *et al*., 2001; Moser *et al*., 2009). In our analysis, CLK-2 is widely conserved in nematodes with some exceptions. In clade I, although the PANTHER (Thomas *et al*., 2022) domain PTHR15830 ‘Telomere length regulation protein Tel2 family member’ is present in all the species, no orthology relationship with *C. elegans* could be identified in the *Trichinellidae*. In contrast, in clade III, both the protein domain and orthology relationship were identified in all the species included in our analysis. In clade IV, both the domain and orthology relationship could be identified in the *Meloidogyne* species but Clk-2 appears to be totally absent from the whole *Aphelenchoididae* and only sparse evidence was found in *Strongyloididae*, similarly to MRT-2. In clade V, candidate clk-2 orthologs could be identified in all the species included in our analysis except *Pristionchus pacificus* although the protein domain was present.

### A complex and composite repeat is enriched at the ends of some contigs

*Meloidogyne* genomes lack the canonical telomeric repeats of nematodes and no other simple repeats were found at contig ends. Furthermore, not even orthologs of telomerase or any of the single- or double-stranded telomeric DNA-binding proteins known in *C. elegans* were found in these species. Besides, it has been observed that ALT-mediated telomere lengthening was associated with notable variations in telomeric sequences (Lee *et al*., 2014). Therefore, we hypothesized other sequences possibly more complex than perfect 6-12mers repeats and not multiplied by telomerases might be present at chromosome ends and play the role of telomeric DNA. Thus, we searched for enriched motifs at contig ends, allowing degenerated imperfect repeated sequences. In *M. incognita*, this led to the identification of a composite repeat made of three different 80bp motifs, ranked by e-value significance as follows. The first one, that we call motif-1, was almost non-degenerated and the two others, called motif-2 and motif-3 were mostly made of stretches of 4 to 6 Gs separated by a few more (motif-3) or less (motif-2) degenerate nucleotides (Figure 2, Supplementary Fig. 7). The composite repeat unit was constituted of motif-2-motif-3-motif-1, in this order, interspersed by non-conserved spacer sequences. Retrieving and aligning all the repeat units present at the contig extremities as well as the spacer sequences allowed deducing a ca. 250-300 bp consensus sequence depending on inclusion or exclusion of the most degenerated (N) positions (Supplementary Fig. 8).

**Figure 2:**
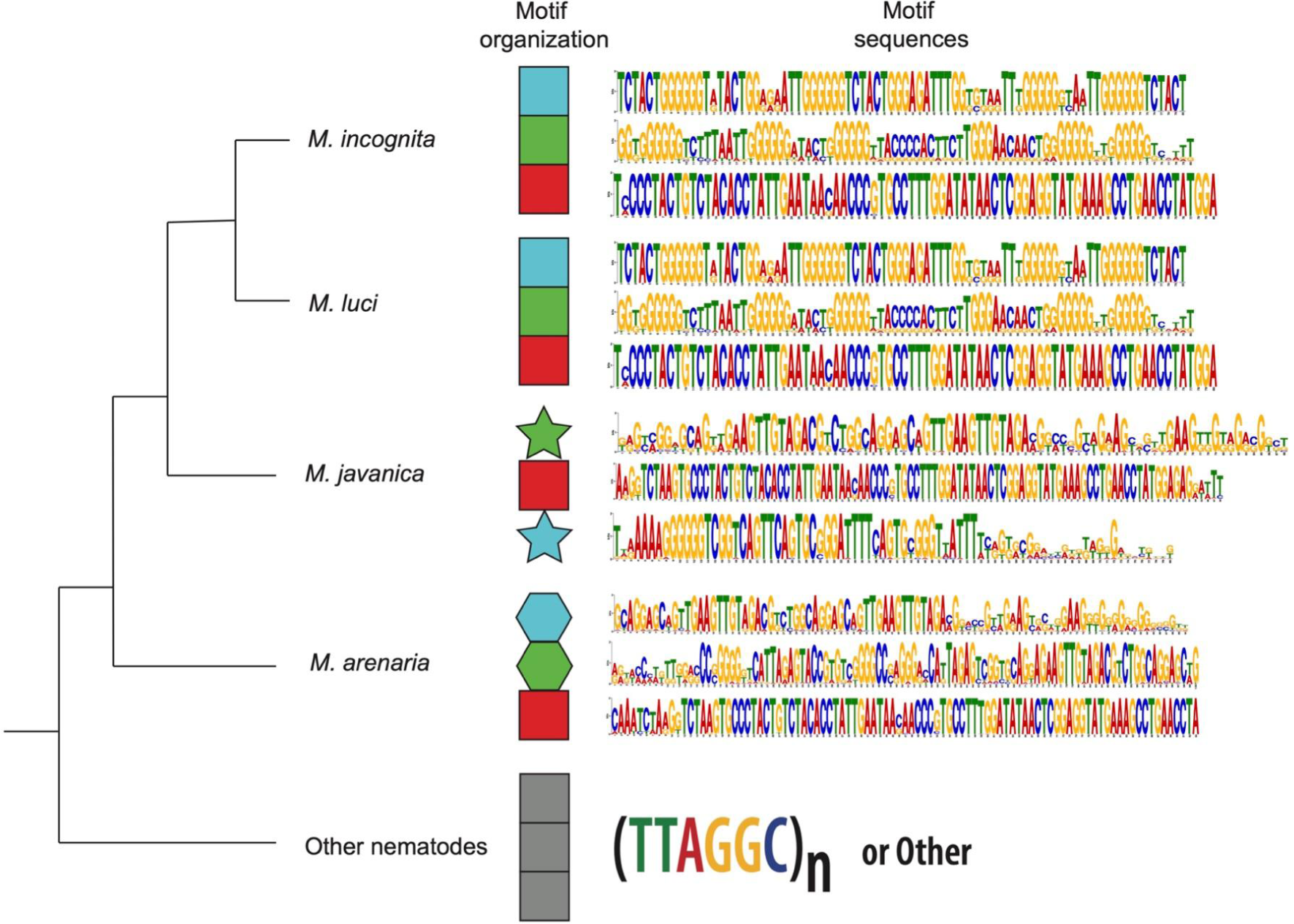
Organization and sequence of enriched motifs at *Meloidogyne* contig ends. Each color on the motif organization indicates a different motif. The red motif (motif 1) having the highest e-value is not degenerated and conserved among *M. incognita*, *M. luci*, *M. arenaria*, *M. javanica*. The G-rich and more degenerated Motifs 2 and 3 are in blue and green, respectively, with different forms representing their non-conservation across *Meloidogyne* species. The motif sequence and conservation within each species is represented in the motif sequences column. Each color in the logo represents a different nucleotide, and their size is proportional to the conservation across all the matches.

This consensus sequence was present in 58 contigs of the *M. incognita* genome and formed repeat arrays on 57 of them (Figure 3, Supplementary Fig. 9, Supplementary Table 3). Repeated arrays contained 8 to 187 repeat units and all contained at least motifs 1 and 2 while 49 contained the 3 motifs. These repeat arrays spanned regions ranging from 2.3 to 59.3 kb and were present exclusively either at the beginning or at the end for 51 of the 57 contigs (∼88%). For one large contig (#10) the arrays started 35kb away from the beginning and 5 shorter contigs ranging in length from 2.4 to 43.8 kb were entirely made of these repeat arrays. Thus, excluding those contigs, the repeat arrays were present at the beginning of 25 contigs and at the end of 26 other contigs but never present at both extremities. We noted that, when present at the beginning of contigs, the consensus sequence was repeated on the + strand (22/25 contigs) while when present at the end, it was repeated on the reverse complement (- strand). We checked in the genome annotation whether some protein-coding genes were predicted in regions containing repeat arrays. In most cases (45/57) no gene was predicted within or overlapping terminal repeat array regions. However, in 12 cases, one single gene was predicted, usually upstream of the repeat array regions. These 12 genes did not belong to the same multigene family but were split across several orthogroups with orthologs identifiable in *M. luci*, *M. javanica* and *M. arenaria*.

**Figure 3:**
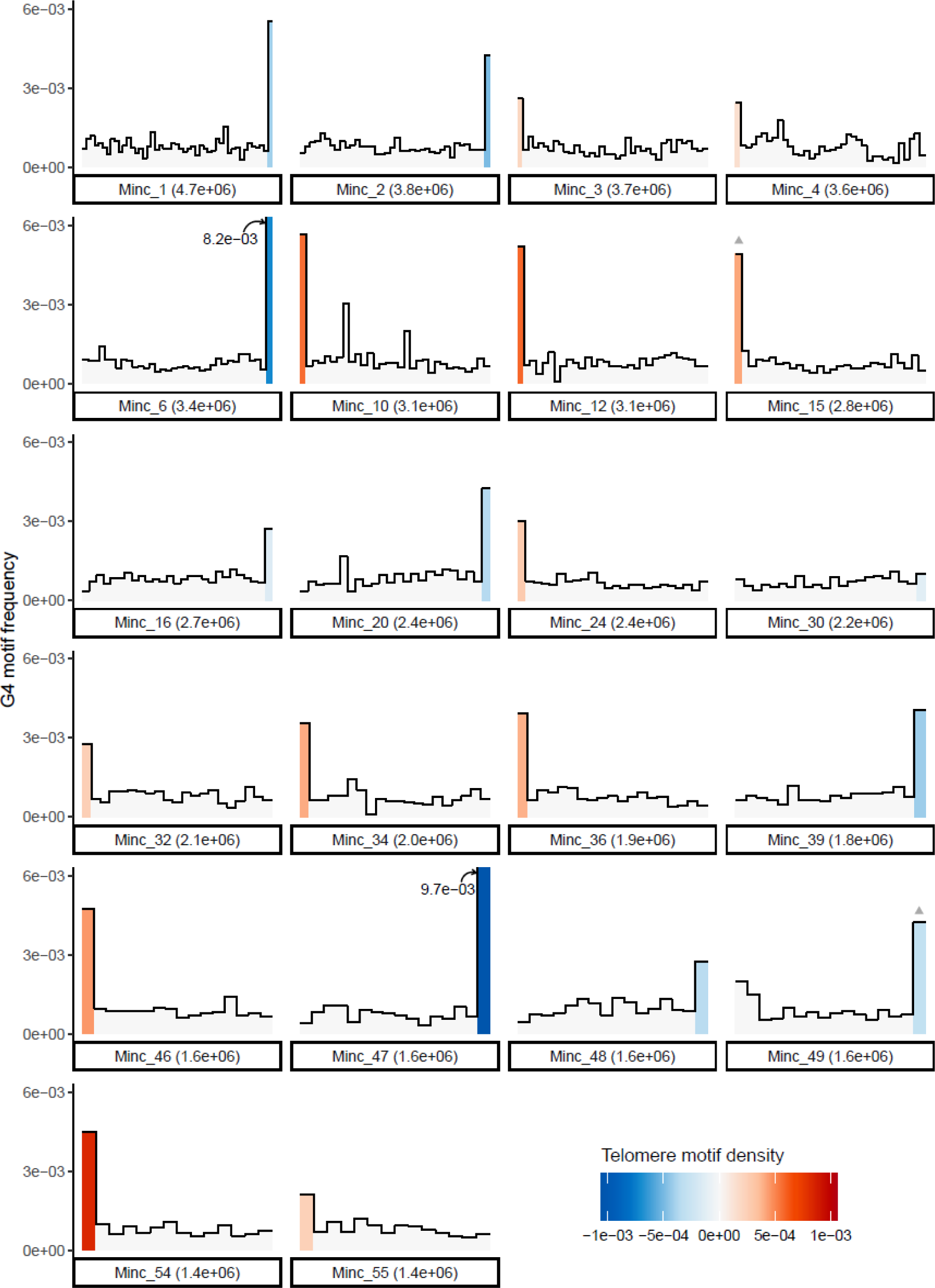
Density of *Minc* composite repeats and G4-quadruplex along genome contigs. The density of *Minc* composite repeat per window is represented by a color gradient with positive values (red) indicating a density in the sense strand and negative values (blue) indicating a density on the reverse complement strand. Gray triangles above bars indicate regions of the contigs where less than 100% of the repeats are on the same strand. The heights of bars in the histogram represent the density of G4-quadruplex forming regions in the same windows. Only contigs containing repeat arrays and bigger than the N80 value are represented, the same information for the rest of repeat-containing contigs is available in (Supplementary Fig. 9).

Because no contig displayed this complex composite repeat at both ends, we investigated whether another motif could be present at the other ends. First, we selected all contigs with a length >1Mb and already containing arrays of the composite repeat at one end. Using the same strategy as explained in methods, we searched for enriched motifs in the first or last 2kb at the opposite end. No evident motif could be identified as enriched, repeated and conserved at the other end of the bigger contigs. We extended this search to all contig beginnings or ends that were not already composed of the complex repeat but again, no evident enriched repeated motif could be found.

We investigated whether the complex repeat identified in *M. incognita* was conserved in the other *Meloidogyne* genomes, including those assembled in this study. Searching the consensus sequence of the *M. incognita* repeat against the *M. javanica* and *M. arenaria* genome assemblies showed that although the consensus sequence matched those genomes, the matches were mostly partial and restricted to the terminal region of the consensus which contains the non-degenerated motif-1 (red). In the *M. arenaria* genome, the *Minc* repeat matched 2,045 positions and was repeated in arrays of three to 125 repeats on 60 different contigs (Supplementary Table 4). Similarly to *M. incognita* motif-1 was repeated either at the beginning or at the end of the contigs with the exception of two contigs that were entirely made of the repeats and two others that contained a repeat array at one end and another at 60kb and 1Mb, respectively, from the other ends. When repeated at the beginning of contigs, motif-1 was generally on the + strand while it was generally on the - strand when repeated at the end of contigs, like in *M. incognita*. Searching specifically for motif-2 and motif-3 revealed, respectively only 2 and 4 hits in the same contig (#109) of *M. arenaria* in a region that also contains motif-1 repeated 9 times at ∼263 kb from the end. In *M. javanica*, the *Minc* repeat formed arrays in 57 contigs, with 31 arrays exclusively at the beginning, 19 exclusively at the end (Supplementary Table 5). Unlike in *Minc* and *Mare*, two *Mjav* contigs (6 and 11) displayed repeat arrays at both ends. Three other contigs had repeat arrays at more than 100kb from the end and two others had repeat arrays in the middle of the contig. We noted that the arrays were limited to a maximum of 21 repeat units while in *M. incognita* and *M. arenaria* the number of repeats can surpass 100. All the repeat arrays present at the beginning of contigs were in the + strand and all those at the end were on the - strand, including for the two contigs containing arrays at both extremities. A search for *Minc* motif-2 and motif-3 returned respectively two and three hits in *Mjav*, all in the same contig and in a region that already contained a repetitive array of motif-1 at 131 kb from the end of contig 147.

Overall, only motif-1 of the *Minc* composite repeat seemed to be conserved in *Mjav* and *Mare* and formed repeated arrays at one extremity of multiple contigs. Therefore, we investigated whether another, slightly different repeat formed arrays at contig ends in *Mjav* and *Mare*, using the same search strategy as originally used in *Minc*. We were able to identify three distinct repeated motifs for each species, present in 60 and 67 contigs, respectively (Figure 2, Supplementary Figures 10A and 11A). One of these motifs was, as expected, the equivalent of the non-degenerated *Minc* motif-1 (red), while two other motifs were different from *Minc* motifs 2 and 3 and also not conserved between *Mjav* and *Mare* (Figure 2). Although the *Mjav* and *Mare* motifs 2 and 3 were relatively G-rich, their consensus sequences were more degenerated than those of *Minc* and were not repeated in the same order as in the *Minc* composite repeat (Figure 2). Using the *Mjav* and *Mare* consensus repeat sequences, we found repeat arrays in the same contigs and the same regions as previously, except that these new consensus sequences fill the gaps left by the absence of match from *Minc* motifs 2 and 3.

#### The complex terminal repeats are specific to polyploid mitotic parthenogenetic root-knot nematodes

Besides the genomes sequenced in this study we also investigated whether the composite repeats found in *Minc*, *Mjav* and *Mare* as well as the motifs that constitute them matched other nematode genomes. We searched in publicly accessible nematode genome assemblies that had a N50 length of at least 100 kb, including those from *M. chitwoodi*, *M. exigua*, *M. luci*, and *M. graminicola*. We found that the *Minc* composite repeat was conserved (including the 3 constitutive motifs) in the publicly available genome assembly of *M. luci* (Susič *et al*., 2020). Like in the other species, it was mostly present repeated at either contig beginnings in the + strand or at contig ends in the - strand, and the repeat unit was constituted of motif-2, motif-3 and motif-1 like in *M. incognita* (Supplementary Table 6). This observation is consistent with a recent phylogenomic analysis of the nematoda phylum suggesting a phylogenetic position of *M. luci* closer to *M. incognita* than to the other polyploid mitotic parthenogenetic root-knot nematodes (Ahmed *et al*., 2022). However, none of the composite repeats or constitutive motifs found in the polyploid mitotic parthenogenetic root-knot nematodes were conserved in the other *Meloidogyne* species analyzed, that are all diploid meiotic species. The composite repeats and their constitutive motifs were also absent from the rest of nematode genomes investigated, suggesting they are specific to the mitotic parthenogenetic *Meloidogyne* species.

More broadly, a search against the NCBI’s nt library with the *Minc* repeat as query returned one single significant hit. Interestingly, this hit was against an *M. incognita* sequence (accession # S68778.1) described, as early as 1991, as a species-specific sequence that could be used as a diagnostic tool (Chacon *et al*., 1991). The match was with a region of the repeat that contains part of motif-3 and the whole motif-1. As could be expected, consensus repeated sequences of *M. arenaria* and *M. javanica* also returned this sole hit against the NCBI’s nt library and only on a region corresponding to the non-degenerated motif-1.

Therefore, besides the clade constituted by polyploid and mitotic parthenogenetic root-knot nematodes, the complex composite repeat identified at contig ends in *M. incognita*, *M. arenaria* and *M. javanica* are not conserved in any other nematode genomes or any other species represented in the NCBI’s nt library so far.

#### No evidence that the terminal repeats of *Meloidogyne* contigs are transposable elements

*Drosophila* and other *Diptera* insects that have lost telomerase have replaced simple telomeric repeats by retro-transposons. Furthermore, previous analyses have suggested transposable elements are active in *M. incognita* (Kozlowski *et al*., 2021). Therefore, we investigated whether the composite repeats present at some contig extremities in mitotic parthenogenetic *Meloidogyne* could be related to transposable elements as well.

We used the EDTA software (Ou *et al*., 2019) to predict and annotate transposable and other repetitive elements in the *Minc, Mjav* and *Mare* genomes (Supplementary Table 7). We checked whether some repetitive elements were predicted in the regions containing the composite repeats we had identified. In *Minc*, EDTA predicted two repetitive sequences in these regions, one described as unclassified repetitive regions and the second one annotated as a putative CACTA TIR transposon (MITE/DTC code). We checked whether these annotated repeats contained a combination of the three motifs constituting the composite 2-3-1 *Minc* repeat. The alignment of each of the three motifs against the EDTA repeat sequences, showed that the motifs 3 and 2 were present in the repetitive sequence TE00000264, unassigned to a known (retro)transposon family, while the motif 1 corresponds to the repetitive sequence annotated as a CACTA TIR DNA transposon (TE00000823). We found this annotation as a DNA transposon surprising as transposable elements described so far as playing the role of telomeres are rather retro-transposons. We further investigated this case and searched whether additional evidence would support this repeat as being a CACTA DNA transposon. No evidence for a transposase or any of the protein-coding genes usually found in CACTA DNA transposon could be identified in this predicted repeat. Furthermore, searching this annotated repeat (TE00000823) against the Rebase28 database (Bao *et al*., 2015) returned no significant hit. Hence, this assignment as a CACTA DNA transposon is most likely an annotation error from EDTA.

No evidence for a transposase or any other protein-coding gene could also be found in the *Minc* composite repeat itself, and the sequence returned no hit against Repbase28. The same results were obtained with *Mjav* and *Mare* composite repeats. Therefore, no evidence supports any of the identified terminal repeats could be related to a known transposable element.

#### The terminal repeats of *M. incognita*, *M. javanica* and *M. arenaria* are predicted to form G4-quadruplexes

One characteristic feature of canonical telomeres, conserved across many eukaryotes, is the presence of a G/C bias with a G-rich telomeric repeat in 3’ and a C-rich reverse complement on the other strand (Bryan, 2020). The G-rich telomeric regions are capable of forming four-stranded G-quadruplexes and these secondary structures are assumed to play important roles in the correct functioning of telomeres (Bryan, 2020). It has even been hypothesized that the evolutionary conservation of G-rich telomeric repeats is related to the importance of forming these G4 quadruplexes. G4 quadruplexes have also been associated with ALT-positive cancer cells and are suspected to play a role in the ALT telomere lengthening pathway. Two motifs (2 and 3) of the *Minc* composite repeat display stretches of Gs interspersed by short non G-rich spacers. The *Mjav* and *Mare* repeats present at contig ends also contain G-rich regions albeit less marked than those of *M. incognita* (Figure 2). Therefore we investigated whether segments of the composite repeats of the three Meloidogyne species could form G4-quadruplexes. Regions forming possible G4-quadruplexes secondary structures were predicted on the three *Minc*, *Mjav* and *Mare* composite repeats. The *Minc* repeat was the one with the highest number of predicted G4 structures (103) and this concerned both G-rich motifs 2 and 3. In *Mare* and *Mjav* composite repeats, respectively 21 and 3 regions were predicted to form potential G4-quadruplexes and they were restricted to their G-rich motifs 3. To investigate whether regions forming G4-quadruplexes were a characteristic of the terminal repeats we have identified, we also predicted these secondary structures on the whole genomes. In *Minc*, we could observe a clear enrichment of segments forming G4-quadruplexes in the genomic regions corresponding to the terminal repeat arrays, with the rest of the genome being otherwise poor in G4-forming segments (Figure 3, Supplementary Fig. 9). Similar results were observed in the *Mjav* and *Mare* genomes although the difference between terminal regions containing the repeat arrays and the rest of the contigs was less marked than in *Minc* (Supplementary Figures 10B and 11B).

Overall, the composite repeats at the extremities of several contigs display G-rich stretches and are predicted to form G-quadruplexes. Although these features are reminiscent of known telomeres in eukaryotes, it must be confirmed that these repeats are actually present at chromosome ends and not the result of assembly artifacts.

### The composite repeat is at one single end of *M. incognita* chromosomes and is a candidate telomeric DNA

To determine the actual localization on *M. incognita* chromosomes of the composite repeat found at the extremities of some contigs, we performed fluorescent *in situ* hybridization (FISH). Specific primers were designed on the consensus sequence of the *Minc* composite repeat (Supplementary Fig 8) and encompassed the whole motifs 1 and 2 as well as the end of motif-3. Amplification of the candidate telomeric repeat showed ladder-like organization visible on agarose electrophoresis gel (Supplementary Fig 12) that is typical for satellite DNA. This result reinforces the hypothesis that the Minc composite repeat is unrelated to transposable elements and rather a satellite DNA. We performed FISH experiments on a total of 10 slides of *M. incognita* to localize the repeat on the chromosomes. Although it is difficult to evaluate the precise number of chromosomes due to their large number and small sizes, it can be roughly estimated between 45-47. This estimation is consistent with a previous analysis on a different stain of *M. incognita* which counted 46 chromosomes (Despot-Slade *et al*., 2021) and with an initial meta-analysis of hundreds of populations which showed the majority of populations had between 41 and 48 chromosomes (Triantaphyllou, 1985).

Even though metaphases are very rarely found, we successfully localized the composite repeat identified in the genome assembly on the chromosomes. Indeed, > 50 evaluated nuclei showed that the repeat form arrays at telomeric and subtelomeric regions of almost all chromosomes (Figure 4). We repeatedly observed only one or two chromosomes without signal. Detailed analyses showed a previously unexpected distribution pattern of candidate telomeric repeats exclusively at one end of the chromosomes (Figure 4), in line with our *in silico* analysis. Signal intensity varied between chromosomes, indicating differences in repeat array length of telomeric/subtelomeric sequence.

**Figure 4.**
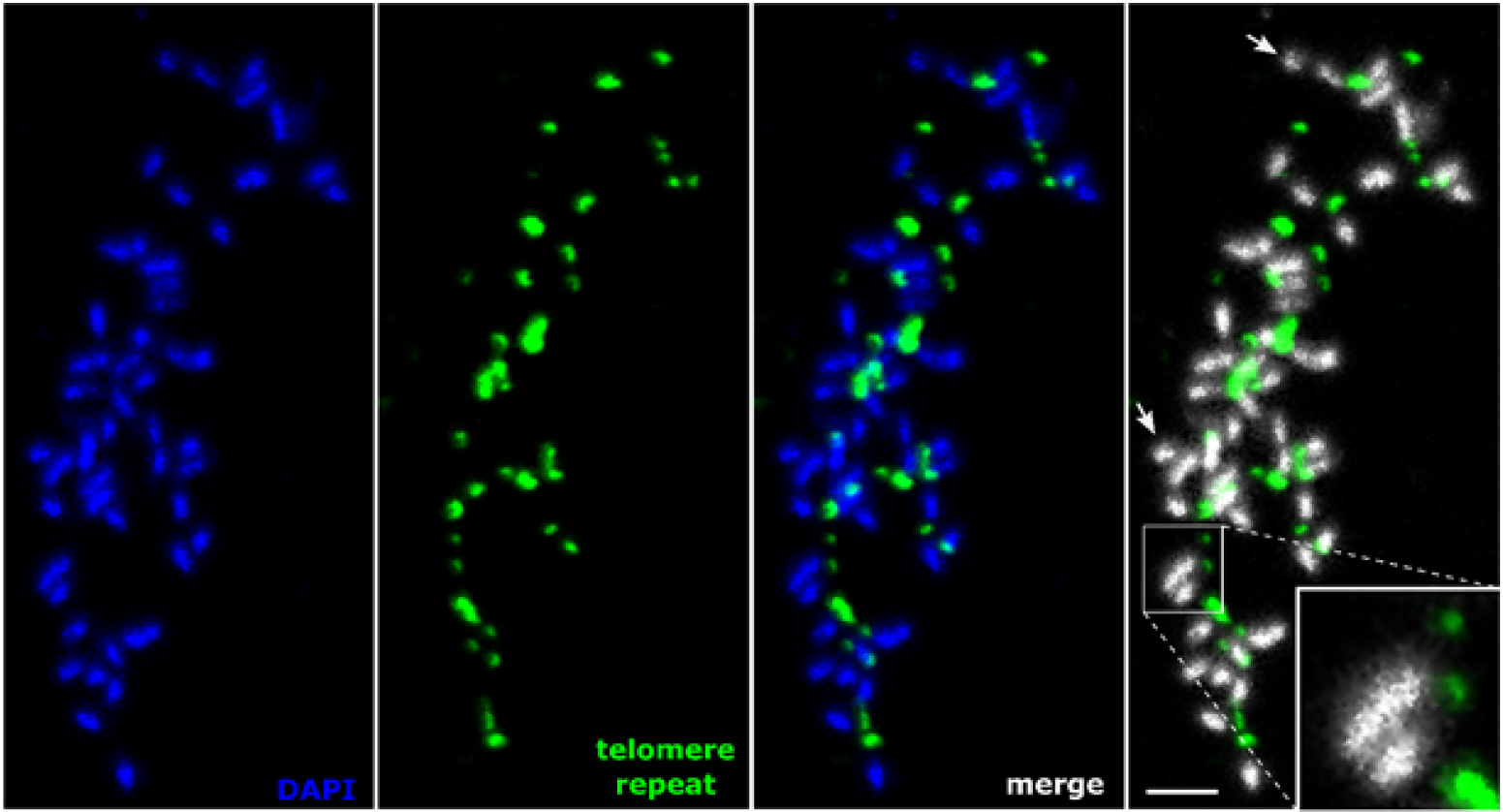
Telomeric chromosomal localization of the *Minc* composite repeat using fluorescence in situ hybridization in metaphase. Specific primers are used for amplification and labeling of the *Minc* composite repeat that was identified at the end of several contigs in the genome assembly. Chromosomes are counterstained with DAPI, arrows represent chromosomes with no visible telomere signal at the end, scale bar = 2 µm.

To exclude the possibility that the presence of one telomeric region per chromosome is a result of U-shaped chromosomes with overlapped telomeric sequences we further investigated the telomeric signals in prophase (Figure 5).

**Figure 5.**
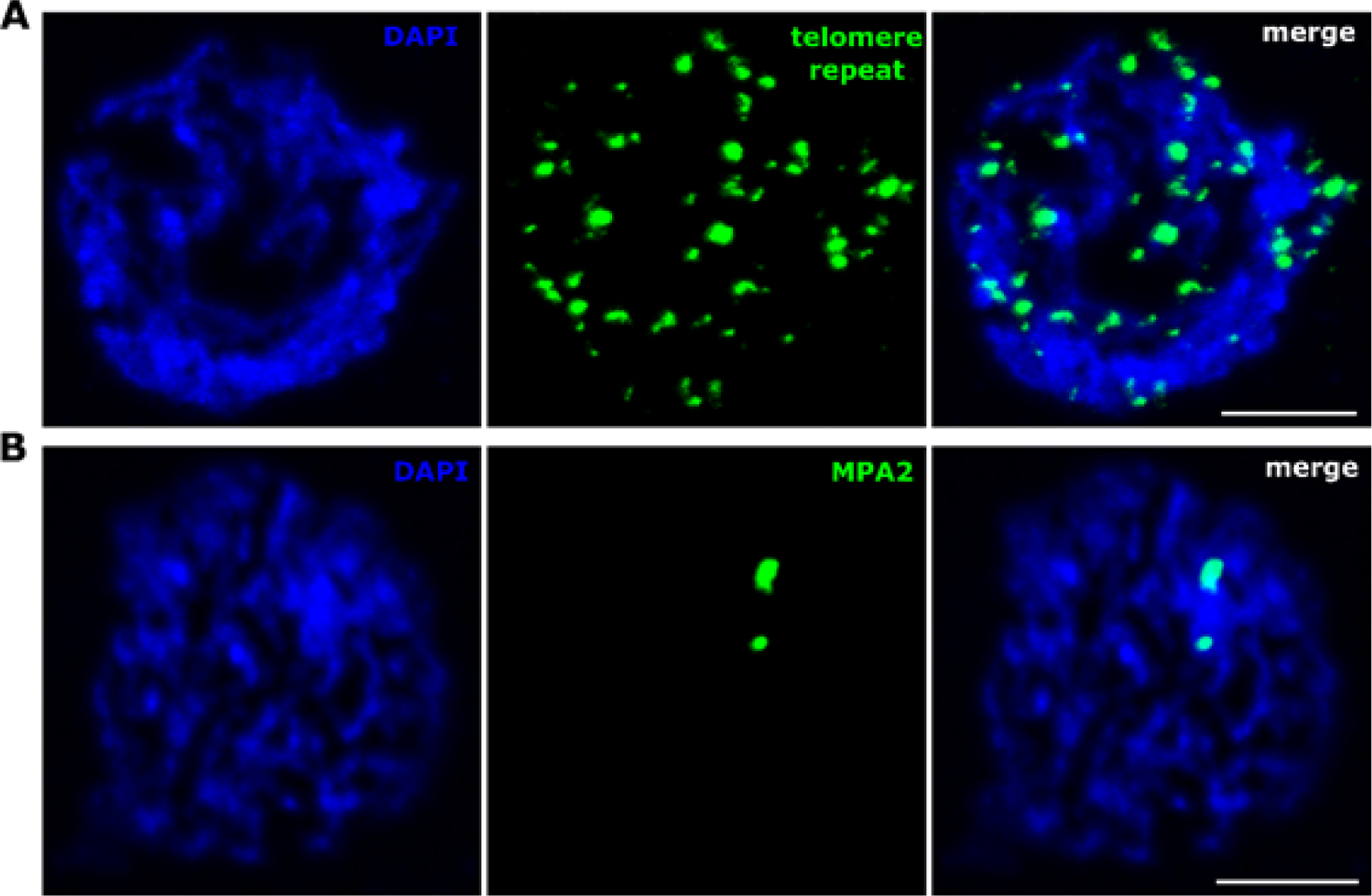
Localization of the Minc candidate telomeric repeat in prophase nuclei. A Distribution of labeled *Minc* repeat sequence. B Labeled probe specific for MPA2 (Meštrović *et al*., 2006) sequence and used as a positive control for FISH. Chromosomes are counterstained with DAPI, scale bar = 5 µm.

Comparison of signals in both prophase and metaphase showed a similar pattern with about 40-45 signals which support the hypothesis that a composite telomeric repeat is exclusively located at one end of most *Minc* chromosomes. These observations are consistent with genome assembly data which showed the composite repeat was exclusively present either at the beginning or at the end of ca. 50 contigs and never at the two ends.

Overall these results confirm the composite repeat identified at one extremity of 49 contigs has a telomeric localization at one end of almost all chromosomes and most likely constitutes a new kind of species-specific telomeric sequence in *M. incognita*.

### The telomeric repeated sequence is transcribed in *M. incognita*

TElomeric Repeat-containing RNA (TERRA) results from the transcription of telomeric regions whether they are constituted of simple repeats like in most eukaryotes, or more complex repeats like transposons in *Drosophila melanogaster* (Azzalin *et al*., 2007; Kordyukova *et al*., 2018). These transcribed telomeric regions constitute a common feature of eukaryotic genomes and they are suspected to play diverse important regulatory roles, including in humans (Bettin *et al*., 2019; Revy *et al*., 2023). For instance, TERRA in mammals is involved in the formation of RNA–DNA hybrid structures, known as R-loop, which play a role in the maintenance of telomeres in ATL positive cells. We investigated whether evidence for transcription of the candidate telomeric repeats could be found in *M. incognita*, for which numerous transcriptomic resources are available. Using previously published transcriptome data for the same isolate of *M. incognita* (Blanc-Mathieu *et al*., 2017), we assembled *de novo* the transcriptomes of four different developmental life stages: eggs, pre-parasitic second stage juveniles (J2), a mixture of late J2, J3 and J4 parasitic stages (J3-J4) and adult females. We then searched for the presence of the three motifs constitutive of the candidate complex telomeric repeats in the assembled transcriptomes. The motifs could be identified in 65 predicted isoforms from 14 assembled transcripts, suggesting the candidate telomeric repeats are transcribed (Table 3, Supplementary Table 8). Two isoforms of a transcript in the J2 stage transcriptome contained both motif-1 and motif-2. In the egg stage, one transcript had two isoforms containing both motif-2 and motif-3 while a third isoform from the same transcript contained motif-1. The rest of the isoforms contained only one of the three motifs. The fact that no single isoform contained the three motifs together might be linked to the difficulty in assembling *de novo* from short read data, sequences containing repetitive low-complexity regions.

**Table 3:**
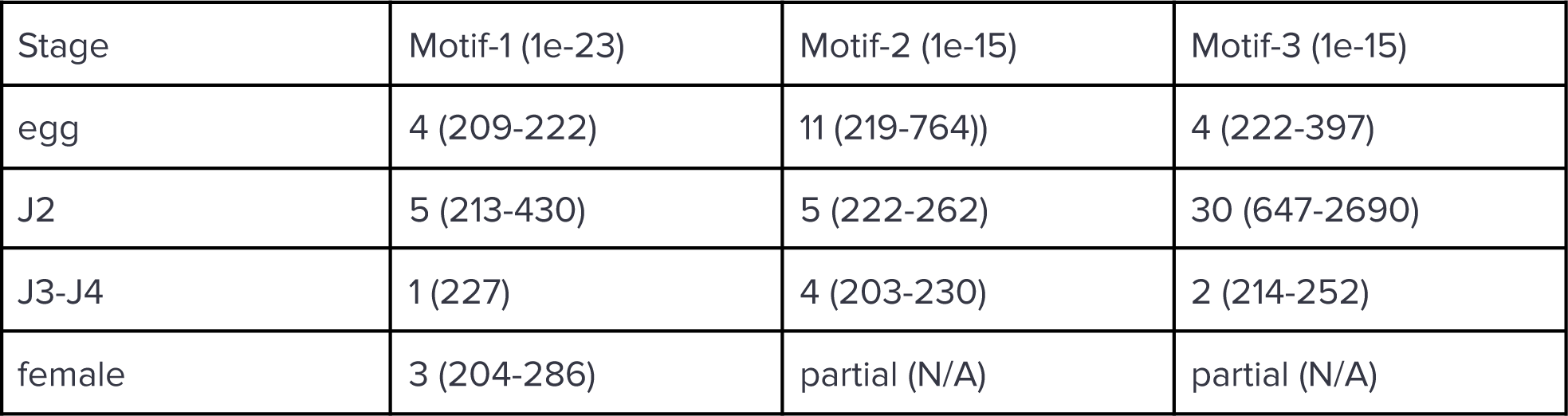
Number of assembled transcripts (including isoforms) containing telomeric motifs and their length distribution in parentheses. Partial means only partial match with the motifs yielding e-value above the threshold.

We found that 9 of the 14 transcripts containing the repeat motifs had at least one isoform that mapped to the *M. incognita* genome assembly with at least 94% identity on at least 80% of their length. Three other transcripts had mapping isoforms but none respected the coverage and identity threshold and two other transcripts had no isoform mapping the genome (Supplementary Table 8).

As short-read transcriptome data could be prone to assembly errors on highly repetitive sequences, we also performed the same suite of analyses on *M. incognita* IsoSeq transcriptome data from mixed developmental stages retrieved from the NCBI (PRJNA787737 SRP350177). We identified 22 iso-seq reads that contained at least one of the three motifs and mapped them to the *M. incognita* genome assembly (Supplementary Table 9). One long (2045 bp) Iso-seq read aligned on its whole length on contig 24 of the *M. incognita* genome. The alignment was on the minus strand at the beginning of the contig in a region spanning two repeated units of the *Minc* telomeric sequence and the terminal region of a protein-coding gene (Minc_v4_shac_contig_24g0207501) just upstream of the repeat array. Another shorter Iso-seq read (293 bp), containing motif-3 in full length as well as motifs 2 and 1 in almost full length, mapped at terminal parts of contigs 1, 79 and 47 in the *M. incognita* genome in regions corresponding to the repeat arrays. Another three reads contained two out of three motifs, and were also mapped to telomeric regions.

Overall evidence from both *de novo* Illumina assembly and Iso-seq data suggest that, similarly to TERRA in vertebrates and other animals, the telomeric repeat of *M. incognita* is transcribed.

## Discussion

Long read sequencing of three allopolyploid root-knot nematodes allowed assembling their genomes at a contiguity level reaching Mb scale and separating the three AAB (*Minc*) and four AABB (*Mjav, Mare*) subgenomes. We identified complex composite repeats mostly present at one extremity of several contigs in the three species and possessing one region (motif-1) conserved across the three species and two other species-specific regions. The repeat possesses several characteristic features of known telomeric DNA in other species such as being G-rich and able to form G4 quadruplexes, being stranded and transcribed with the potential to form G-loops. Using FISH, we confirmed in *M. incognita* that this repeat has a telomeric localization at one extremity of chromosomes and is most likely the telomeric DNA repeat. It can be hypothesized that the repeats identified in *M. luci*, *M. javanica* and *M.arenaria* that share similar features and mostly terminal distribution along the contigs also represent telomeric DNA in these species.

The discovery of these stranded telomeric repeats revealed how chromosomes start (or end) in these *Meloidogyne* species. FISH experiments have confirmed the repeats are at only one extremity of chromosomes. Thus, contigs bearing these repeats can now be placed and oriented accordingly. This pivotal ensemble of information will guide further scaffolding efforts using Hi-C or optical mapping towards full chromosome resolution of these complex genomes. Initial cytogenetics analyses performed on hundreds of populations for these and other *Meloidogyne* species suggested the numbers of chromosomes were mostly (41-46) for *Minc*, (42-48) for *Mjav* and (51-56) for *Mare* (Triantaphyllou, 1985). The number of contigs bearing the composite repeats in the three species are consistent with these early cytogenetics analyses and suggest our assemblies at the contig level constitute a solid base to further scaffold the genomes at end-to-end chromosome scale resolution.

In addition to helping scaffold future versions of the genomes, the discovery of these specific composite telomeric repeats in *Meloidogyne* opens new perspectives for further research. The mechanisms by which these composite repeats might be multiplied at one end of chromosomes remain to be discovered. They cannot be multiplied by a telomerase enzyme because no evidence for this enzyme was identified in any *Meloidogyne* species. Furthermore, unlike in *Drosophila*, these repeats are not retrotransposons and seem to be unrelated to any known transposable elements and more likely constitute a kind of satellite DNA. Therefore, multiplication by (retro)-transposition also appears unlikely. An attractive hypothesis is that they might be maintained and multiplied via an ALT pathway by a recombination-based mechanism such as observed in some human cancer cells and also hypothesized as the ancestral telomere multiplication mechanism in eukaryotes (Bryan, 2020). Consistent with this hypothesis, the telomeric repeats in *Meloidogyne* possess markers of ALT pathway based on their localisation, orientation, the heterogeneity in the repeats and the presence TERRA-like transcript which could lead to G-loops formation, in the absence of telomerase.

Another question is to identify the proteins that bind these unusual telomeric repeats since no ortholog of any of the single-stand or double-strand telomeric DNA binding proteins of *C. elegans* were found in Meloidogyne species. Telomere-binding proteins such as those composing the shelterin complex play essential roles in telomere maintenance, stability and function. In that perspective it is interesting to note that *Drosophila* have replaced the shelterin by another complex playing the same role and composed of four proteins encoded by orphan genes and called terminin (Raffa *et al*., 2011). It is tempting to speculate that in Meloidogyne too, orphan proteins might have been recruited to bind, stabilize and protect the composite telomeric repeats. Alternatively or additionally, G-quadruplexes or G-loops could play a role in the recruitment of proteins involved in the maintenance of the telomere integrity. Another common feature of telomeres is that telomeric regions are transcribed and produced TERRA for (telomeric repeat containing RNA). TERRA then interacts with telomeric DNA and proteins and has been shown to play central regulatory functions in telomeres (Azzalin and Lingner, 2015). In nematodes, it is unknown whether RNA from transcribed telomeric DNA also play important roles. In *M. incognita*, we found that several transcripts contained the composite telomeric repeat and might constitute an analogue of TERRA with functions that remain to be investigated. Based on the orientation and the presence of putative G-quadruplexes, we can hypothesize that these TERRA-like transcripts could be involved in the formation of R-loops or G-loops that could play a role in the ALT pathway developed by Meloidogyne species .

In the future, we would like to further investigate our hypotheses using experiments such as DNA immunoprecipitation technique with an antibody recognising the DNA–RNA hybrids in R-loops and further analyze the TERRA-like transcript to access for potential G-loop formation that could be associated with ALT pathway in this organisms.

Finally, because these composite telomere repeats seem to be specific to these devastating *Meloidogyne* species and because telomeres play functions essential for genome integrity and species survival, this discovery opens new translational perspectives towards more efficient and specific pest control methods targeting telomeres.

## Material and Methods

### *Meloidogyne* samples, DNA extraction, library preparation and sequencing

The same *Minc*, *Mjav* and *Mare* isolates previously sequenced with short reads (Blanc-Mathieu *et al*., 2017), and maintained in the INRAE collection were reared on tomato plants and used for DNA extraction.

A total of 11, 8 and 9 ONT libraries for *Minc*, *Mjav* and *Mare*, respectively, were prepared using the SQK-LSK108 and SQK-LSK109 ligation sequencing kits, following kit changes and Oxford Nanopore instructions over time (Supplementary Table 10). Libraries were loaded on MinION or PromethION R9.4 Flow Cells, according to the Oxford Nanopore protocols, targeting over 100X coverage per genome.

To allow contigs correction, PCR free Illumina libraries were prepared using the Kapa Hyper Prep Kit (KapaBiosystems, Wilmington, MA, USA), following the manufacturer’s instructions. Libraries were then quantified by qPCR using the KAPA Library Quantification Kit for Illumina Libraries (KapaBiosystems), and library profiles were assessed using a High Sensitivity DNA kit on an Agilent Bioanalyzer (Agilent Technologies, Santa Clara, CA, USA). Libraries were then sequenced on an Illumina NovaSeq instrument (Illumina, San Diego, CA, USA), using 250 base-length read chemistry in a paired-end mode, generating approximately 100X coverage for each genome (Supplementary Table 11).

### Genome assembly

#### ONT reads QC and filtering

We first used guppy-v6.0.6 in super high-accuracy mode with a minimal quality score of 7 (--min_qscore 7) to base-call reads from the raw sequencing signal. Raw sequencing libraries and the configuration files used for the basecalling are described in Supplementary Table 10. We then analyzed the reads quality and length distribution as a quality-check using cONTent, a home-made script available at https://github.com/DjampaKozlowski/cONTent.

Following QC, we used nanofilt (De Coster *et al*., 2018) to crop the 50 first and the 30 last nucleotides as these regions showed reduced per-base quality. We then selected reads that were at least 2 kb long and had a quality score Q > 12. For *M. incognita*, this returned 1,709,063 higher quality cleaned reads out of a starting number of 6,619,891 raw reads. The same method was used for both *M. arenaria* and *M. javanica*, which before the filter had 10,506,836 and 11,506,089 raw reads, respectively. After filtering we obtained 2,592,069 reads for *M. arenaria* and 3,144,046 for *M. javanica*.

#### Illumina reads QC and filtering

Low-quality nucleotides (*Q*<20) from both ends of the reads that passed Illumina filtering were discarded. Illumina sequencing adapters and primer sequences were removed from the reads. Then, reads shorter than 30 nucleotides after trimming were discarded. These trimming and removal steps were achieved using in-house-designed software based on the FastX package (https://github.com/institut-de-genomique/fastxtend). The last step identifies and discards read pairs that are mapped to the phage phiX genome, using SOAP aligner (Li *et al*., 2008) and the Enterobacteria phage PhiX174 reference sequence (GenBank: NC_001422.1). This processing, described in (Alberti *et al*., 2017), resulted in high-quality data. After all these steps, reads shorter than 167 bp were eliminated.

#### Assembly

We assembled the genomes from ONT cleaned data using NECAT (Chen *et al*., 2021) with default parameters and an input genome size of 200 Mb for *M. incognita*, 310 Mb for *M. arenaria* and 300 Mb for *M. javanica*. These input genome sizes do not influence final assembly size but are used to sample the longest highest quality long read covering the genome at at least 40X. After *M. incognita* assembly and bridging, NECAT formed 298 contigs for a genome size of 199,975,050 bp and a N50 length of 1,856,931 (longest contig = 4,672,579 bp, L50#= 37). For *M. arenaria* we obtained 398 contigs with an assembly size of 305,105,593 bp and a N50 of 2,046,842. For *M. javanica* we obtained 371 contigs with an assembly size of 298,215,187 bp and a N50 of 2,079,353 (Table 4).

**Table 4:** raw statistics of genome assembly based on ONT long reads.

| Species | raw reads | clean reads | reads used by Necat | contigs after polishing | contigs w/o mitochond. | genome size (Mb) |
| --- | --- | --- | --- | --- | --- | --- |
| *M. incognita* | 6,619,891 | 1,709,063 | 287,593 | 293 | 291 | 199.42 |
| *M. arenaria* | 10,506,836 | 2,592,069 | 447,241 | 378 | 377 | 304.33 |
| *M. javanica* | 11,506,089 | 3,144,046 | 279,109 | 365 | 364 | 297.81 |

#### Polishing and contamination check

For polishing we performed three different steps for each species. Using the assembly produced by NECAT, we ran two rounds of Racon (Vaser *et al*., 2017) using the cleaned ONT reads. The resulting fasta file from the first mapping was used as a reference genome for the second round of mapping and correction. For each species, between 6 (*M. javanica*) and 20 (*M. arenaria*) contigs which did not have enough coverage by the long reads, were dropped. The new set of polished contigs from Racon were further polished by one round of Medaka (https://github.com/nanoporetech/medaka). Finally, the contigs polished with long reads were further polished with cleaned PCR-free Illumina short reads using one round of Hapo-G (Aury and Istace, 2021) with default parameters. This led to polished assemblies with 293, 378 and 365 contigs for *M. incognita*, *M. arenaria* and *M. javanica*, respectively. We then use blobtools to detect possible contaminations based on contig GC content, coverage and taxonomic assignment (Kumar *et al*., 2013). The analysis revealed no contamination but allowed identifying contigs that corresponded to the mitochondrial genomes (Supplementary Figure 4). These contigs were removed from the final assembly, and the final number of contigs was 291, 377 and 364 for *M. incognita*, *M. arenaria* and *M. javanica*, respectively (Table 4).

### Genome ploidy, divergence and completeness assessment

We used JellyFish (Marçais and Kingsford, 2011) with the default k-mer size of 21 to enumerate k-mers on the cleaned short reads for each species. Using the counts generated by JellyFish we generated histograms using the jellyfish histo command and adjusted the -h parameter to allow k-mer repeated up to 1 million times to make sure downstream analysis will not be limited by the default maximum allowed occurrence of k-mers. We then used SmudgePlot (Ranallo-Benavidez *et al*., 2020) to infer the ploidy level for each species (Supplementary Figure 1). Finally, we used GenomeScope2 (Ranallo-Benavidez *et al*., 2020) with the previously identified ploidy levels and generated k-mer histograms to estimate genome sizes and average percent of nucleotide divergence between homoeologous genome copies (Supplementary Figure 2).

We estimated genome completeness using two strategies. First, we used KAT (Mapleson *et al*., 2017) to estimate how far the diversity of k-mers present in the cleaned short reads was retrieved in the genome assemblies for each species (Supplementary Figure 3). Second, we used CEGMA (Parra *et al*., 2007) and BUSCO (Manni *et al*., 2021) to estimate which percentage of evolutionarily conserved single-copy genes could be retrieved in the genome assemblies. We used CEGMA version 2.5 with default parameters to identify eukaryotic evolutionarily conserved genes and their average numbers in the *Meloidogyne* genomes. We used BUSCO version 5.4.4 with the Metazoa odb9 dataset and default parameters to estimate the percentage of widely conserved animal genes retrieved in complete and partial length in the three genomes.

### Gene predictions and duplications detection and classification

Predictions of gene models in *M. incognita*, *M. javanica*, *M. arenaria* and genomes were done with the fully automated pipeline EuGene-EP version 1.6.5 (Sallet *et al*., 2019). EuGene has been configured to integrate similarities with known proteins of *Caenorhabditis elegans* (PRJNA13758) downloaded from Wormbase ParaSite (Howe *et al*., 2017) as well as the “nematoda” section of UniProtKB/Swiss-Prot library (UniProt Consortium, 2018), with the prior exclusion of proteins that were similar to those present in RepBase (Bao *et al*., 2015). We used as transcriptional evidence, transcriptome data for *M. incognita*, as it is the *Meloidogyne* species with the most comprehensive expression data available. RNA-seq data from pre-parasitic J2, J2-J3 and adult female stages (Blanc-Mathieu *et al*., 2017) were assembled *de novo* using Trinity (Haas *et al*., 2013) followed by a cleanup that retains for each trinity locus only the transcript that gives the longest ORF. The dataset of *M. incognita* assembled transcriptome was aligned on the genomes of the four *Meloidogyne* species using Gmap (Wu and Watanabe, 2005) and except for *M. incognita* the option “cross-species” was used. Only alignments spanning 30% of the transcript length with at least 97% identity were retained. Statistics of gene predictions in the four species are available in Supplementary Table 1.

The EuGene default configuration was edited to set the “preserve” parameter to 1 for all datasets, the “gmap_intron_filter” parameter to 1, the minimum intron length to 35 bp, and to allow the non-canonical donor splice site “GC”. Finally, the Nematode specific Weight Array Method matrices were used to score the splice sites (available at this URL: http://eugene.toulouse.inra.fr/Downloads/WAM_nematodes_20171017.tar.gz).

To identify duplicated genomic regions we used McScanX (Wang *et al*., 2012) with default parameters. For each species, we used as input to McScanX an all-against-all BLASTp comparison of the proteins and the position of the protein-coding genes in the genome.

### Transposons and other repetitive elements annotation

We predicted transposable and other repetitive elements on the three genome assemblies using EDTA (Ou *et al*., 2019) version 2.1. To allow a more comprehensive annotation of repetitive elements, we used the –sensitive parameters, which uses the RepeatModeler (Flynn *et al*., 2020) annotation to identify other repeats which were not identified previously. We configured RepeatModeler to identify the remaining repeats (sensitive 1), to perform a whole genome repeat annotation (anno 1), and to evaluate the consistency of the previous annotation (evaluate 1), the rest of the parameters were left to default.

### Distribution of telomeric repeats and telomere-associated proteins in nematode genomes

#### Dataset of nematode genomes and proteomes

We collected nematode genome assemblies with a minimal N50 length of 100 kb either at the scaffold or contig level and with predicted protein-coding genes available. We started by the list of sequenced genomes at WormBase ParaSite (Howe *et al*., 2017) and within a given nematode genus we kept up to three genomes with the highest N50 lengths. Then, we complemented this list with the ensemble of nematode genomes available at the NCBI, using the same criteria for selection (N50 length >100 kb, availability of predicted genes and up to three genomes in a given genus). Our selection included the *C. elegans* genome and proteome and we did not apply a limitation of 3 species for the *Meloidogyne* genus, since this was the focal point of our study. Because no gene predictions were available for *Meloidogyne luci* (Susič *et al*., 2020) despite its genome satisfying the N50 threshold, we predicted gene models using the same strategy as used for *M. incognita*, *M. javanica* and *M. arenaria* with Eugene (Sallet *et al*., 2019) and with the *M. incognita* transcriptome as a source of evidence. To anchor our analysis we also included the genome assembly and predicted proteins of the tardigrade *Hypsibius exemplaris* (Yoshida *et al*., 2017) as an outgroup. This resulted in 68 nematode species and one outgroup (Supplementary Table 2).

#### Detection of nematode homologs of *C. elegans* telomere-associated proteins

Starting from *C. elegans* proteins known to be associated to telomeres, including the telomerase Trt-1 as well as telomere-binding proteins Pot-1, Pot-2, Pot-3, Mrt-1, Tebp-1/Dtn-1 and Tebp-2/Dtn-2, we searched for homologs in the other nematode proteomes and genomes selected above. In addition to these proteins we also included Clk-2 and Mrt-2 which have suspected roles in telomere maintenance but are also important DNA damage checkpoint proteins. We employed the following strategy:

i. we retrieved protein motifs characteristic of the *C. elegans* proteins under consideration and used hmmsearch from HMMER3 (Eddy, 2011; Mistry *et al*., 2013) against the predicted proteomes of all the collected species to check for their presence. We set an e-value threshold allowing retrieval in the *C. elegans* proteome of the query protein and no unrelated proteins. For Trt-1, we used the ‘telomerase reverse-transcriptase’ PANTHER (Thomas *et al*., 2022) domain (PTHR12066) with an e-value of 1E-15. Pot-1 presented no specific protein-domain and this domain-based strategy could not be used for this protein. The single-strand telomeric DNA-binding proteins Pot-2, Pot-3 as well as Mrt-1 all possess a ‘ssDNA-binding domain of telomere protection protein’ Pfam (Finn *et al*., 2016) domain (PF16686) which was used as a query with an e-value of 1E-15. For Mrt-2, we used the ‘Repair protein Rad1/Rec1/Rad17’ Pfam domain (PF02144) as a query with an e-value threshold of 1E-40. For the double-strand telomeric DNA-binding proteins Tebp-1 and Tebp-2, we used as query the Panther domain ‘GA binding and activating and spk (spk) domain containing-related’ (PTHR38627) with an e-value threshold of 1E-10. Finally, for Clk-2, we used the Panther domain PTHR15830 ‘Telomere length regulation protein Tel2 family member’ with an e-value threshold of 1e-15.
ii. we performed an Orthofinder analysis of all the proteomes (68 nematodes + 1 tardigrade) to classify the different proteins in orthogroups. We used the option -M msa to calculate multiple sequence alignments and phylogenies for all the orthogroups and the species tree. Then, for each *C. elegans* protein of interest we retrieved the corresponding orthogroup and checked which other species were present in the same orthogroup and thus had a putative ortholog. For the specific case of telomerase we also used the *Ascaris suum* enzyme as a query, because strong evidence for an active telomerase in this species has been reported (Estrem and Wang, 2023).
iii. in case of absence of evidence for a homolog or disagreement between (i) and (ii) we retrieved the *C. elegans* protein sequence for the gene of interest and aligned it in against the genome of the nematode species translated in the 6 frames using tBLASTn with an e-value threshold of 1E-5. We also repeated the same tBLASTn search against WormBase Parasite (Howe *et al*., 2017) to extend the search to different versions of a genome assembly for a given species.

For each species, homologs of *C. elegans* telomere-associated genes were considered possibly present if evidence from (i) and (ii) could be identified. In case of disagreement between (i) and (ii), but significant tBLASTn matches in (iii), homologs were considered present but misannotated. If no evidence from either (i), (ii) or (iii) could be identified, the gene was considered as having no ortholog in the considered species. If only evidence for either (i) or (ii) were present without confirmation by (iii), we considered the presence of an ortholog unsure.

#### Detection of canonical nematode telomere repeats

The 68 nematode genomes were scanned for the canonical (TTAGGC)n nematode telomere repeat using the Telomere Identification toolKit Tidk (https://github.com/tolkit/telomeric-identifier). We used the command tidk find with -c nematoda option and a window size of 1000 nucleotides. The *C. elegans* telomere repeat was considered present if repeated at least 50 times within the 2000 last or first nucleotides of at least 3 contigs. Because repetitive regions might be misassembled in the genomes we also used fuzznuc from the EMBOSS package (Rice *et al*., 2000) to further search (TTAGGC)_2_ dimers anywhere on the genome assemblies. We considered the telomeric repeat possibly present if repeated at least 100 times (50 consecutive dimers). Finally, the absence of the canonical (TTAGGC)n repeat may also be due to repeat filtering during the genome assembly process. Therefore, the canonical telomeric repeat was searched in raw Illumina reads whenever available in SRA. For each species, when available, we chose the SRA data produced with illumina NovaSeq for WGS. We used (TTAGGC)_10_ as a query to perform a blastn search on NCBI, using the SRA library corresponding to the species of interest as a subject and deselecting the low complexity regions filter. We considered the telomeric repeat as possibly present when more than 100 matches of (TTAGGC)_10_ could be retrieved in the reads. The canonical repeat was considered present in a species whenever evidence from either tidk or from fuzznuc plus blast against SRA could be retrieved. When evidence from fuzznuc was found but no SRA data was available the canonical repeat was considered possibly present. In all other cases, it was considered absent.

### Identification of new candidate telomeric repeats

Because the canonical (TTAGGC)n telomeric motif was not found in the *Meloidgyne* genomes, we also searched for clusters of any perfect repeats of sizes 6-12 using the equicktandem and etandem commands from the EMBOSS package (Rice *et al*., 2000).

Because no simple tandem repeats of size 6-12 were identified at contig ends, we searched for more complex and possibly degenerated motifs. For each species, we extracted the first and last 2kb of all contigs and searched for frequent motifs using MEME-suite (Bailey *et al*., 2015) in classic mode on these sequences. Any number of repetitions (anr) mode was selected with iteratively searching motifs from size 50 to 100 with up to 5 different motifs reported. After examination of MAST results from the MEME suite, we realized that two motifs frequently co-occurred and covered almost the whole length of many contig beginnings or ends. We then only selected those contig beginnings and ends that contained repeats of these two motifs and re-launched a new motif search. This new motif search was with max. 3 motifs with a size range of 80-160. The results of MAST allowed identifying for each species (Minc, Mjav and Mare) a suite of 3 enriched motifs with one motif being conserved in the three species.

For each species we considered the suite of 3 motifs as a repeat unit, and extracted all these repeat units in the corresponding genomic regions. We aligned these repeat units using MAFFT v7 (Katoh and Standley, 2013) with option G-INS-i. The obtained multiple sequence alignment was then used as an input to the EMBOSS (Rice *et al*., 2000) Cons program to build a consensus.

We also retrieved for each enriched motif of each species, the corresponding FASTA sequences from MEME results, built a multiple alignment with MAFFT using the same parameters as for the whole repeat unit and constructed a hmm profile for each motif, using hmm-build from the HMMER package (Mistry *et al*., 2013).

### Distribution of the newly identified candidate telomeric repeats in *Meloidogyne* and other genomes

To investigate how the *Minc* composite repeat was distributed in the genome, we used it as a query in a blastn (Camacho *et al*., 2009) search against the genome with an e-value threshold of 1e-35 and no dust filter. Because two of the three motifs and thus the repeat units were different in *Mjav* and *Mare* as compared to *Minc*, we also used the *Mjav* and *Mare* consensus sequences as queries, to repeat the same search in the *Mjav* and *Mare* genomes, respectively.

To investigate whether the three consensus sequences of the repeat units were conserved in other nematodes, we also searched with blastn and with the same parameters in all the other nematode genomes we downloaded in this study (Supplementary Table 2). Furthermore, for each of the three complex repeat units, we also performed an online blastn search against all the nematode genomes present in Wormbase ParaSite (Howe *et al*., 2017) with an e-value of 0.01 and no low complexity filter.

Finally, we also investigated more widely whether the candidate complex telomeric repeats had homologs against the NCBI’s nt library using blastn online with an e-value of 0.01 and the low complexity filter deactivated.

Besides the whole repeat unit, we also investigated how each individual motif constituting them was conserved across *Meloidogyne* and other species. The hmm profile of each motif was searched against the 68 nematode genomes we collected using nhmmer from HMMER (Mistry *et al*., 2013). For the non-degenerated motifs-1 of the three species, we used an e-value of 1e-23 while for the more degenerated motifs 2 and 3, we used an e-value of 1e-18.

### Confirmation of telomeric position of the identified repeats with FISH

Specific primers 5’-TCGTCCATAGGTTCAGGCTT-3’ and 5’-GGGTCTACTTGGGTGGGG-3’ (Supplementary Fig. 8) were designed on the *Minc* consensus candidate telomere sequence using Primer 3 software (Untergasser *et al*., 2007). Probe for FISH experiment was prepared by PCR labeling with biotin-16-dUTP (Jena BioScience) using *M. incognita* genomic DNA as template. PCR reaction was prepared in 25 µL volume containing 1x Colorless GoTaq Flexi Buffer, 2.5 mM MgCl2, 0.1 mM dNTPs, 0.4 µM primers, 1.25 U of GoTaq DNA Polymerase (Promega) and 0.02 ng of gDNA. For PCR settings 3 min at 95 °C of initial denaturation was followed by 35 amplification cycles (20 s at 95 °C, 20 s at 62 °C and 40 s at 72 °C) and 5 min at 72 °C for final extension. Prepared probe was cleaned using QIAquick PCR Purification Kit (Qiagen) and eluted in 30 µL of nuclease-free water and checked on 1% agarose gel for successful biotin incorporation (Supplementary Fig. 12). The corresponding labeled probe had length (200 bp) ideal for hybridization steps during *in situ* localization.

Microscopic slides and subsequent fluorescent in situ hybridization (FISH) was performed as described in (Despot-Slade *et al*., 2021). Briefly, cell suspension from isolated females infecting tomato roots was applied on slides using Cytospin 4 cytocentrifuge (Shandon, ThermoFisher Scientific). Slides were incubated for 20 min at -20 °C in methanol:acetone (1:1) fixative, dried and carried through the FISH. Specimens are pretreated in 45 % acetic acid, incubated for 30 min at 37 °C with RNase A and fixed with 1% formaldehyde in PBS with 50 mM MgCl2. Chromosome denaturation was performed in 70% formamide in 2xSSC at 70 °C for 2 min. On each slide 100 ng of lyophilized labeled probe resuspended in hybridization buffer and denatured at 75 °C for 5 min was applied and incubated at 37 °C overnight. Afterwards, 4x posthybridization washes in 50% formamide in 2xSSC and immunodetection with fluorescein avidin D and biotinylated antiavidin D system (Vector Laboratories) were carried out. Slides were counterstained with DAPI, mounted with Mowiol 4-88 (Sigma-Aldrich) and images recorded using confocal laser scanning microscope Leica TCS SP8 X. For each cytosmear 6-8 z-stacks slices with average thickness of 5 µm are acquired and images were post-processed using ImageJ and Adobe Photoshop software.

#### G4-quadruplex prediction

We used the G4-Hunter software (Brázda *et al*., 2019) with a window size of 25 and a minimal score of 1.2 to detect regions possibly forming G4-quadruplexes on the composite terminal repeats as well as the whole genome sequences of the three *Meloidogyne* species. Using an in-house script, we retrieved only the non-overlapping positions, distant by more than 25 bp, and the score for each G4-quadruplex position, and we plotted their frequency using Rstudio.

#### Transcriptomic support of *Minc* telomeric repeats

We retrieved *M. incognita* RNA-seq Illumina data of four developmental life stages (eggs, pre-parasitic J2, J3-J4 and adult female) from a previous publication (Blanc-Mathieu *et al*., 2017). For each developmental life stage, we pooled the triplicates and performed a de novo transcriptome assembly using Trinity-v2.4.0 (Haas *et al*., 2013). We used nhmmer (Mistry *et al*., 2013) with the hmm profiles of each *Minc* motif as queries to identify assembled transcripts and isoforms containing one or several of these motifs. We repeated the same nhmmer search on an Iso-seq mixed-stage *Minc* transcriptome downloaded from the NCBI (PRJNA787737 SRP350177).

Assembled transcripts as well as Iso-seq reads that contain one or several telomeric motifs were aligned on the *M. incognita* genome using GMAP (Wu and Watanabe, 2005) with default parameters.

## Data availability

All the raw long and short genome sequencing reads for the three species have been deposited in the EBI’s European Nucleotide Archive (ENA) under BioProject PRJEB61149 with detail of each library and accession numbers in Supplementary Tables 10 and 11. Genome assemblies and annotations have been deposited in the French national data repository ‘Recherche Data Gouv’ and will be available upon publication of the paper in a peer-reviewed journal.

## Acknowledgment

We are grateful to the genotoul bioinformatics platform Toulouse Occitanie (Bioinfo Genotoul, https://doi.org/10.15454/1.5572369328961167E12) for providing computing resources. We are grateful to the bioinformatics and genomics platform, BIG, Sophia Antipolis (ISC PlantBIOs, https://doi.org/10.15454/qyey-ar89) for computing and storage resources. We would like to thank Marine Poullet, Julia Truch, Philippe Castagnone-Sereno, Pierre Abad, and Claire Caravel for help, support and stimulating discussions. We are grateful to Paulo Vieira and Sebastian Eves-van den Akker for providing early access to the genome and predicted proteins of *Pratylenchus penetrans*.. This work was supported by the French government, through the UCAJEDI Investments in the Future project managed by the National Research Agency (ANR) under reference number ANR-15-IDEX-01. The authors are grateful to the OPAL infrastructure and the Université Côte d’Azur’s Center for High-Performance Computing for providing resources and support.

