## Supplementary Material for "Unzipped assemblies of polyploid root-knot nematode genomes reveal new kinds of unilateral composite telomeric repeats"

### Supplementary information

#### Supplementary Figure. 1: k-mer estimation of genome ploidy

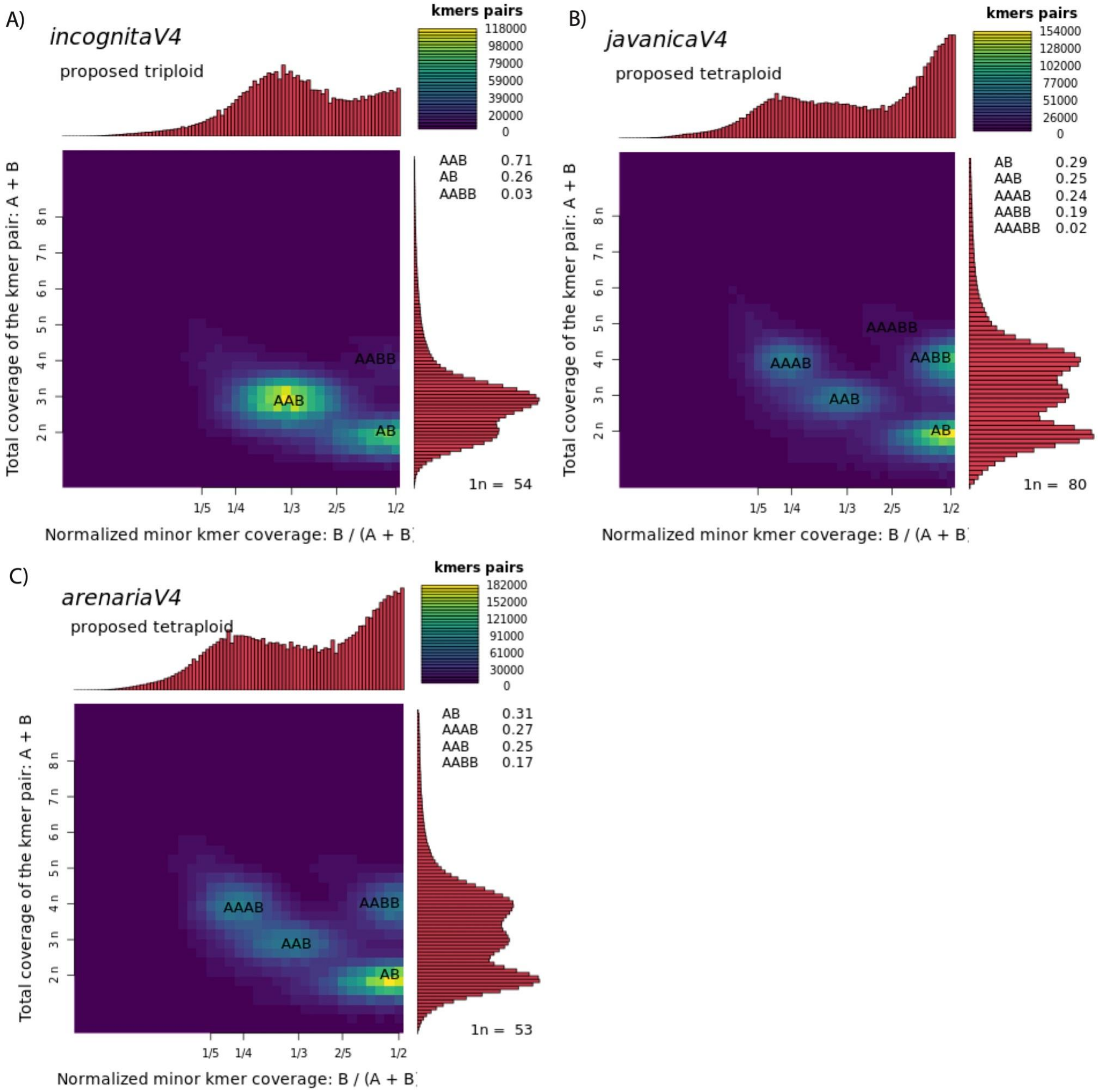

Smudgeplot (Ranallo-Benavidez et al., 2020) k-mer distribution analysis of the Illumina reads used for polishing the contigs predicted a triploid (3n ‘AAB’) genome structure for *M. incognita* (**A**) and tetraploid (4n ‘AABB’) genome structures for both *M. javanica* (**B**) and *M. arenearia* (**C**).

### Supplementary Figure. 2: k-mer estimation of genome size and heterozygosity

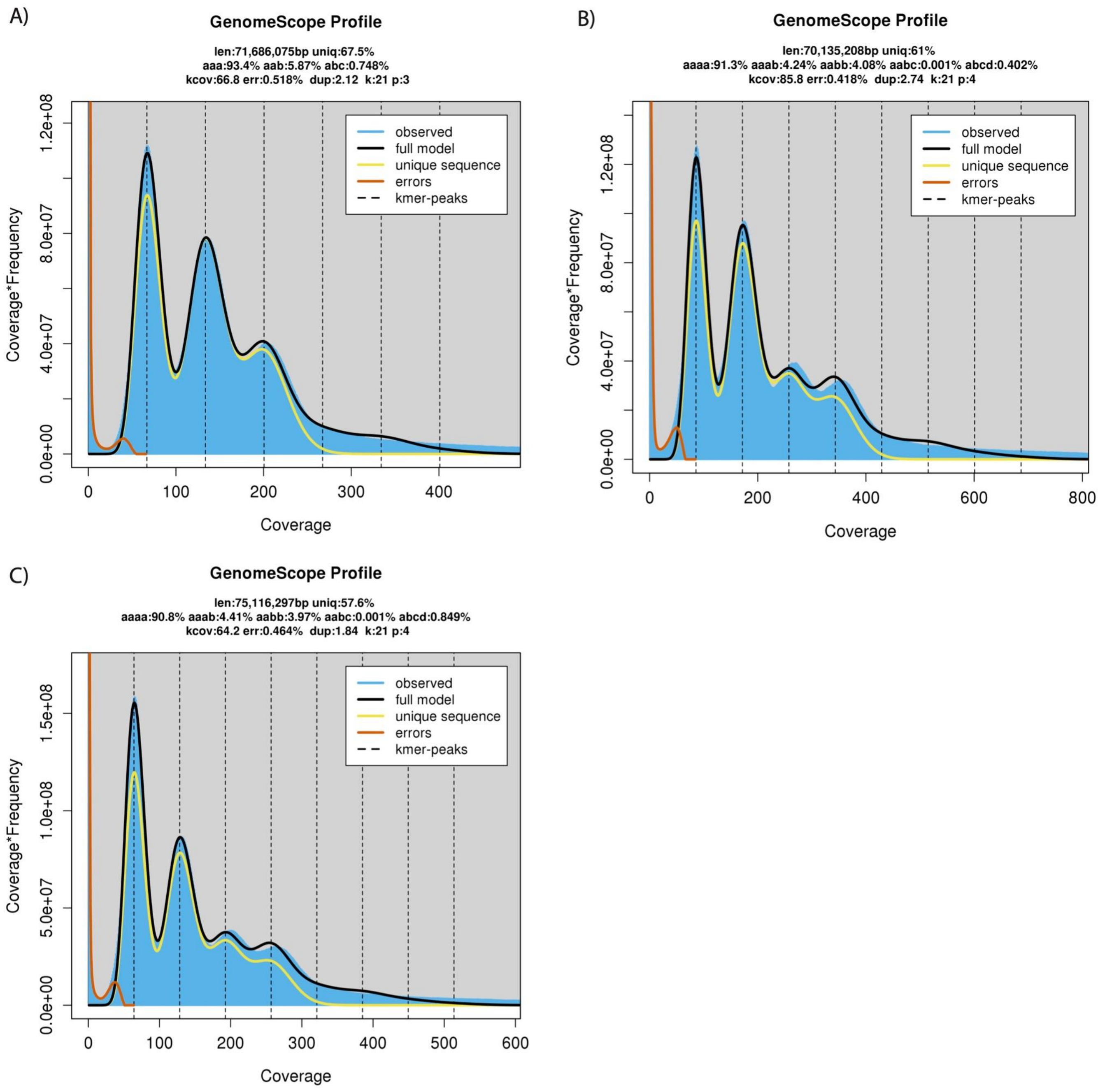

Estimation of genome size and nucleotide divergence between homoeologous genome copies using GenomeScope2 (Ranallo-Benavidez et al., 2020). (**A**) Setting the ploidy at 3n in *M. incognita* yielded an estimated haploid genome size of ca. 71.69Mb (and thus a total genome size of ca 215Mb), with an average nucleotide divergence between homoeologous genome copies of 6.6%. (**B**) Setting the ploidy at 4n in *M. javanica* yielded an estimated haploid genome size of ca. 70.14 Mb (and total genome size of 280.6Mb) with an average nucleotide divergence between genome copies of 8.7%. (**C**) Setting the ploidy at 4n in *M. arenaria* yielded an estimated haploid genome size of ca. 75.12 Mb (total genome size of ca. 300.5 Mb) with an average nucleotide divergence between genome copies of 9.2%.

### Supplementary Figure 3 k-mer estimation of genome assembly completeness

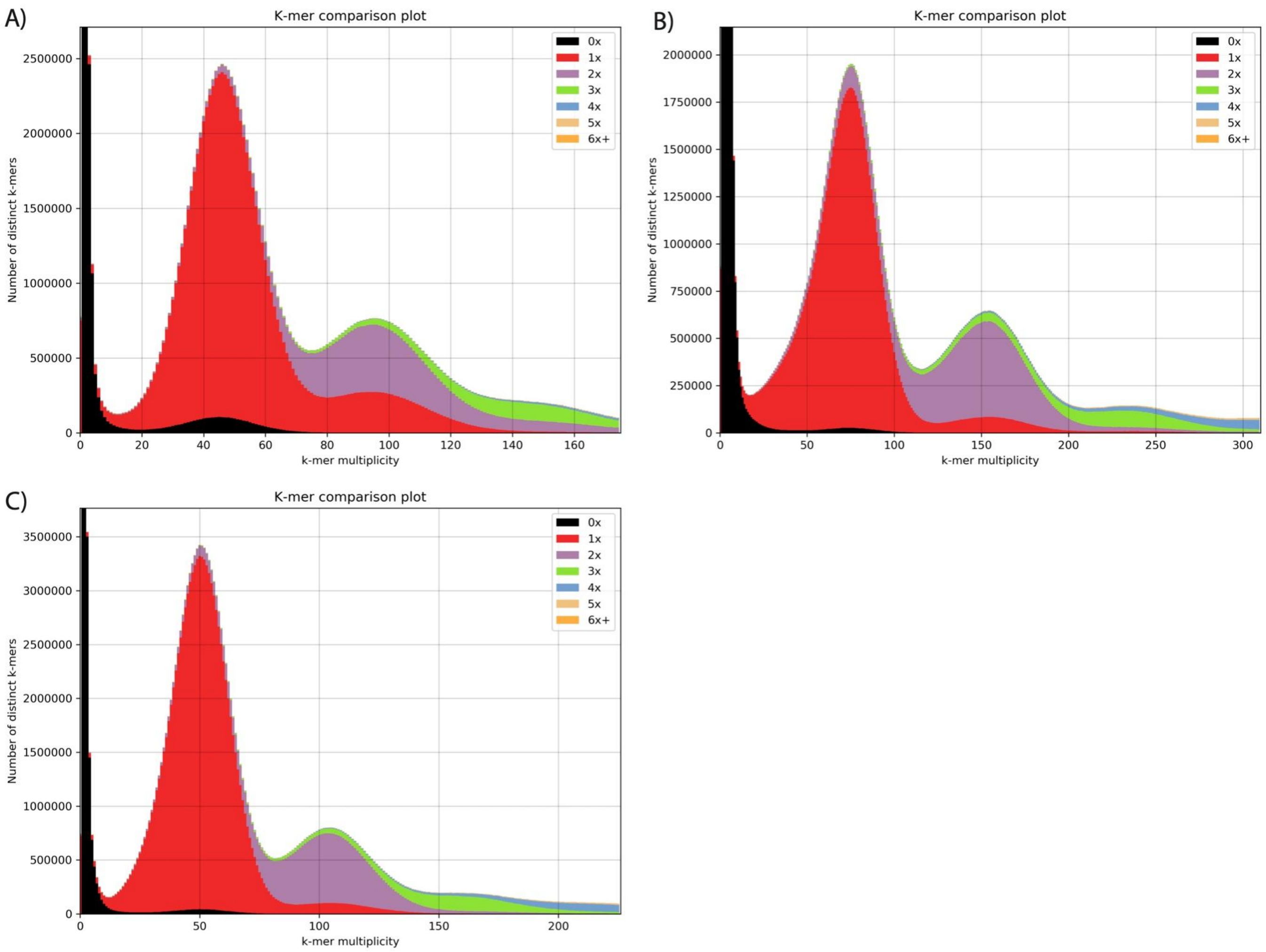

KAT (Mapleson et al., 2017) comparative analysis of the distribution of Illumina reads k-mers in the *M. incognita* (**A**), *M. javanica* (**B**) and *M. arenaria* (**C**) genome assemblies. Very few k-mers in the reads were not represented in the assembly (0X black curve), suggesting almost all the information present in the reads was retrieved in the assemblies. This observation and the congruence between genome assembly sizes and those estimated via flow cytometry (Blanc-Mathieu et al., 2017) suggest complete genomes with homoeologous genome copies having been mostly unzipped during assembly.

### Supplementary Figure 4: genome contamination assessment with blobtools

#### 1) *M. incognita*

A)

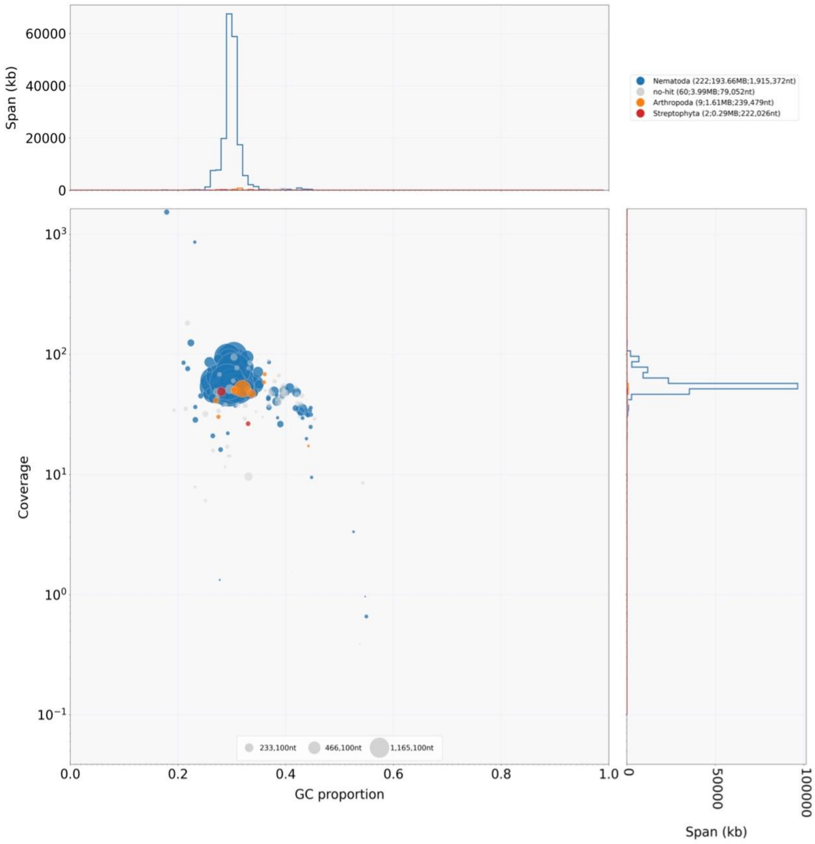

B)

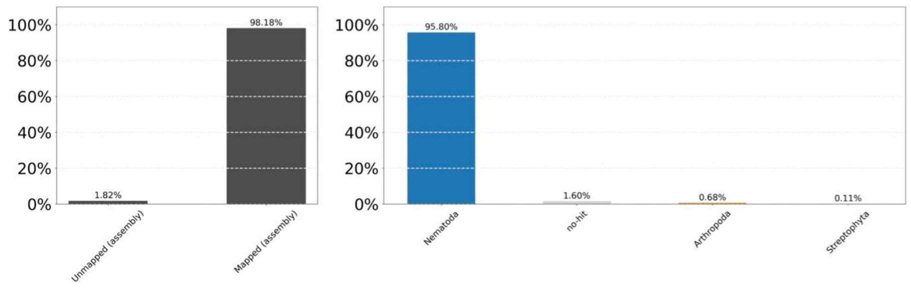

No evident trace of contamination could be identified. Two contigs with very high coverage corresponded to the mitochondrial genome of *M. incognita* and were removed from the final nuclear genome assembly.

### 2) *M. javanica*

A)

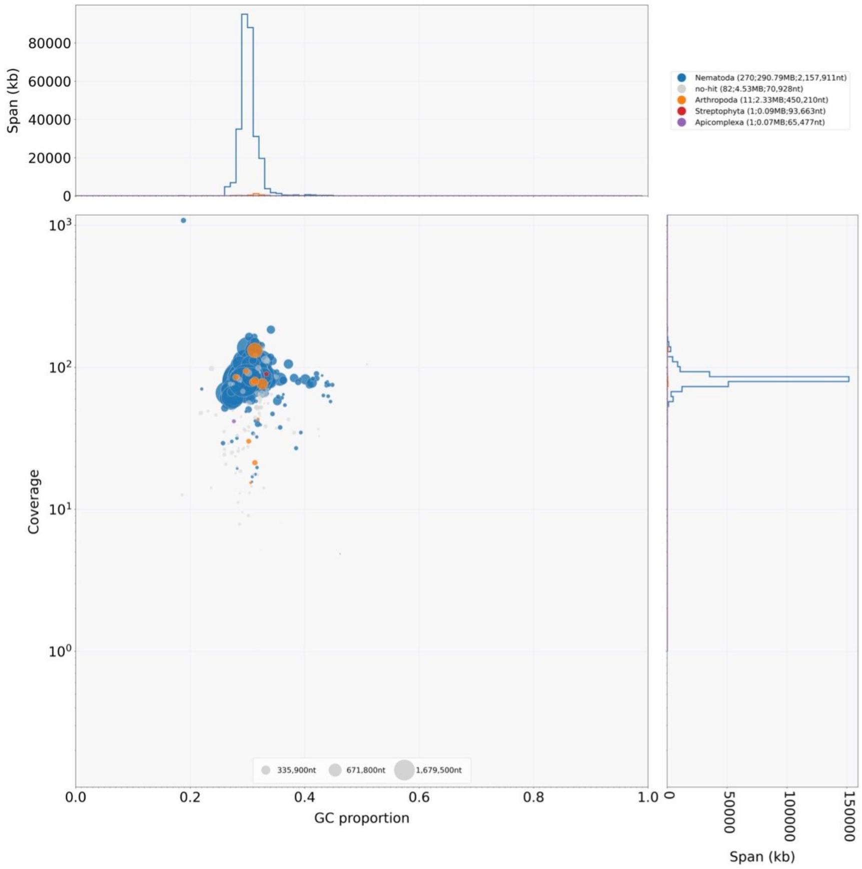

B)

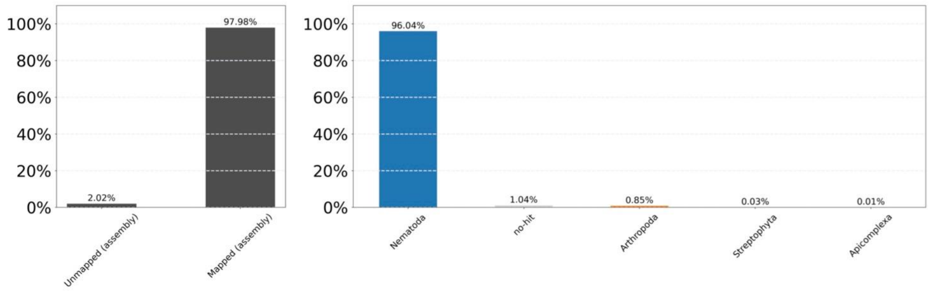

The majority of the contigs had approximately the same coverage, and one contig identified with a high coverage was annotated as the mitochondrial genome of *M. javanica* and removed from the assembly. A low percentage of the contigs was mapped as Arthropoda (0.85%), but this can be the result of lack of information on the NCBI database.

#### 3) *M. arenaria*

A)

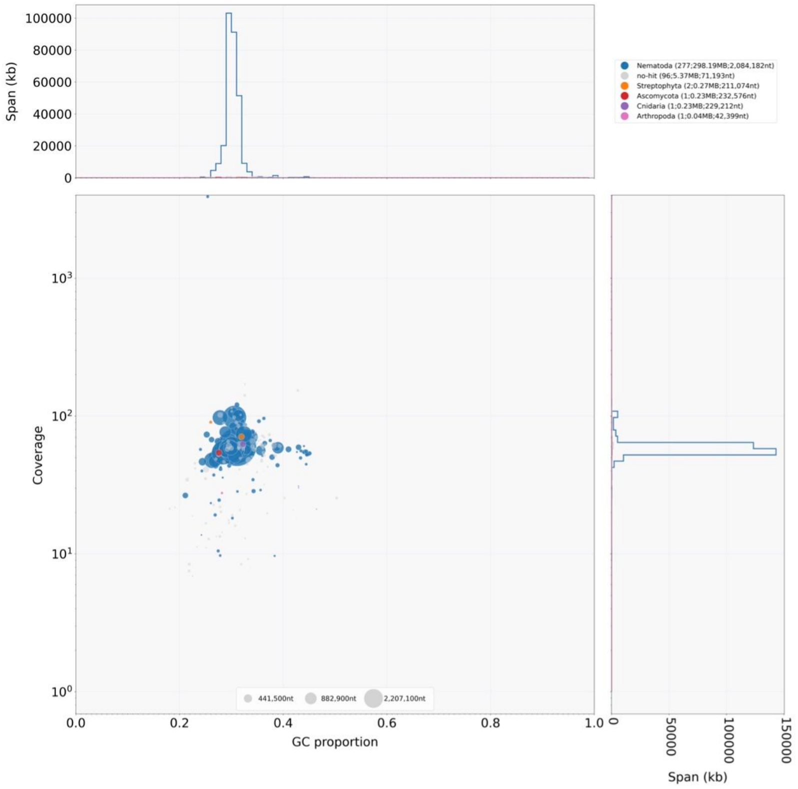

B

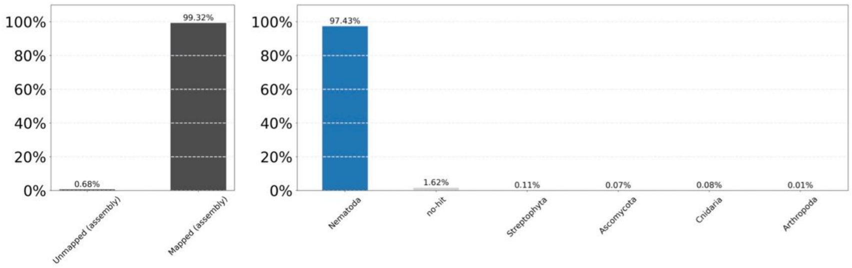

One contig was identified with an high coverage (> $10^3$) compared to the others. This contig corresponded to the mitochondrial genome of *M. arenaria*, and was eliminated from the final assembly. No contamination was observed in the final assembly.

### Supplementary Table 1: Statistics of EuGene predictions

| Feature / species | *Minc* | *Mjav* | *Mare* | *Mluci* |
|---|---|---|---|---|
| Predicted genes | 54,816 | 64,689 | 63,434 | 52,935 |
| Protein-coding | 52,182 | 61,060 | 59,500 | 49,988 |
| Avg. intergenic distance (coding genes) in bp | 1,442.7 | 2,563.3 | 2,745.6 | 1,685.4 |
| Avg. length protein-coding genes (bp) | 2,254 | 2,102 | 2,122 | 2,333 |
| Percent of the genome coding | 23.6% | 19.2% | 18.5% | 22.3% |
| GC% in coding regions | 34.43% | 34.63% | 34.63% | 34.56% |
| Percent of genes with introns | 75% | 82% | 83% | 79% |
| Avg. introns / gene | 4.4 | 4.7 | 4.8 | 4.7 |
| Avg. intron length | 216.4 | 203.8 | 203.1 | 230.6 |
| Percent of noncanonical (GC) splice donor | 1.2% | 1.0% | 1.0% | 1.3% |
| Non protein-coding genes | 2,634 | 3,692 | 3,934 | 2,947 |
| GC% of non-coding genes | 38.6% | 39.97% | 40.9% | 38.3% |
| Percent of genome masked by RED[1] | 15.8% | 23.8% | 24.3% | 16.4% |

1: regions masked by EuGene (Sallet et al., 2019) using RED (Girgis, 2015) prior to gene prediction because they are repetitive with no evidence for transcription.

### Supplementary Table 2: Distribution of (TTAGGC)n repeat and telomere-associated proteins across nematode genomes.

Table available online at:

https://entrepot.recherche.data.gouv.fr/privateurl.xhtml?token=e74d9638-9bc5-4ff8-9ccc-c5348539ceb1

### Supplementary Figure 5: telomeric repeat ancestral state reconstruction

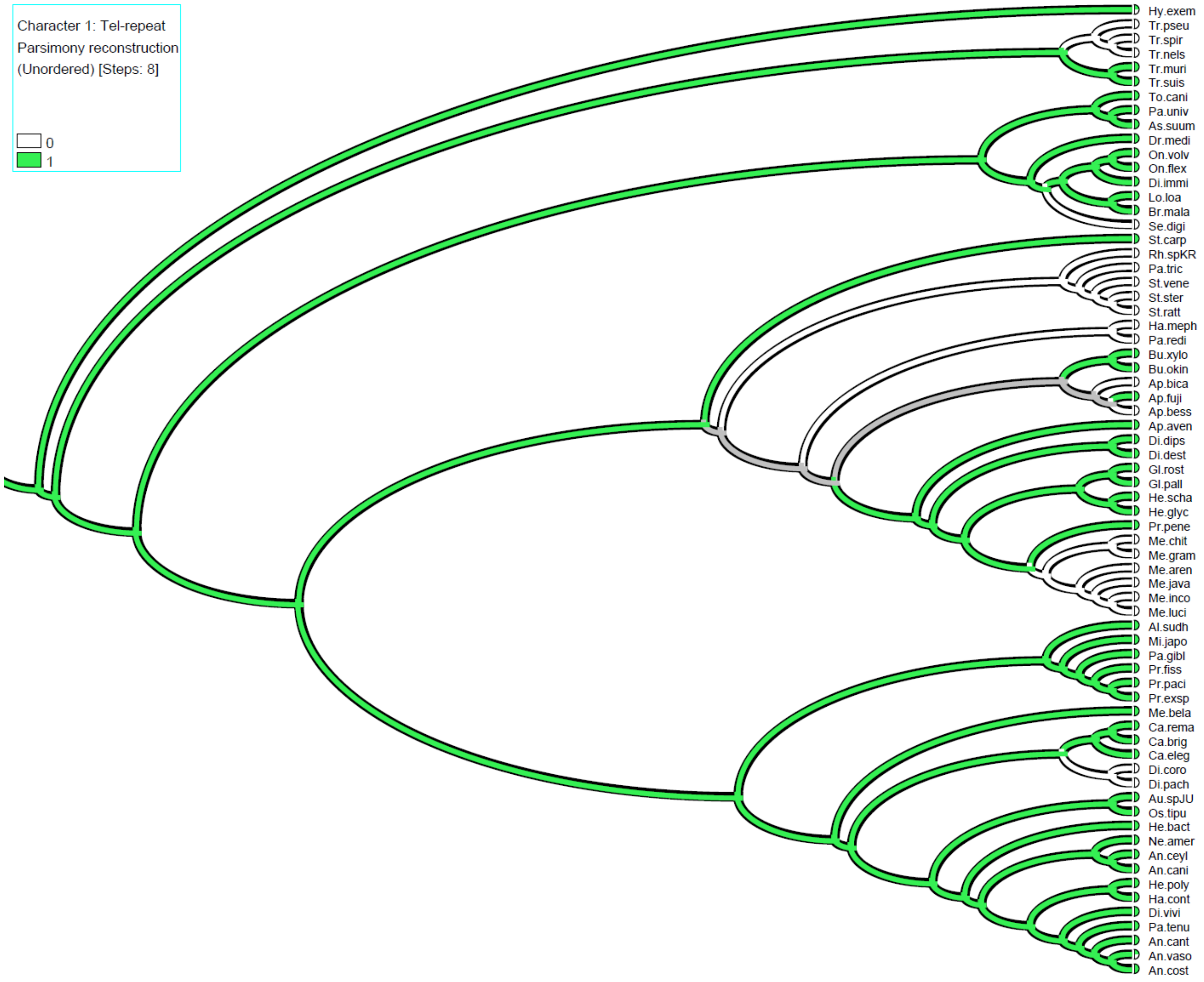

Based on presence / absence of the (TTAGGC)n telomeric repeat of *C. elegans* in nematode genomes, ancestral states were reconstructed using parsimony in Mesquite (Maddison and Maddison, 2014). Green means (TTAGGC)n present and white absent, gray means presence and absence are equi-parsimonious. In the tardigrade *Hypsibus dujardini*, another simple repeat, (GATGGGTTTT)n, was described as a candidate telomeric repeat (Yoshida et al., 2017).

### Supplementary Figure 6: telomerase ancestral state reconstruction

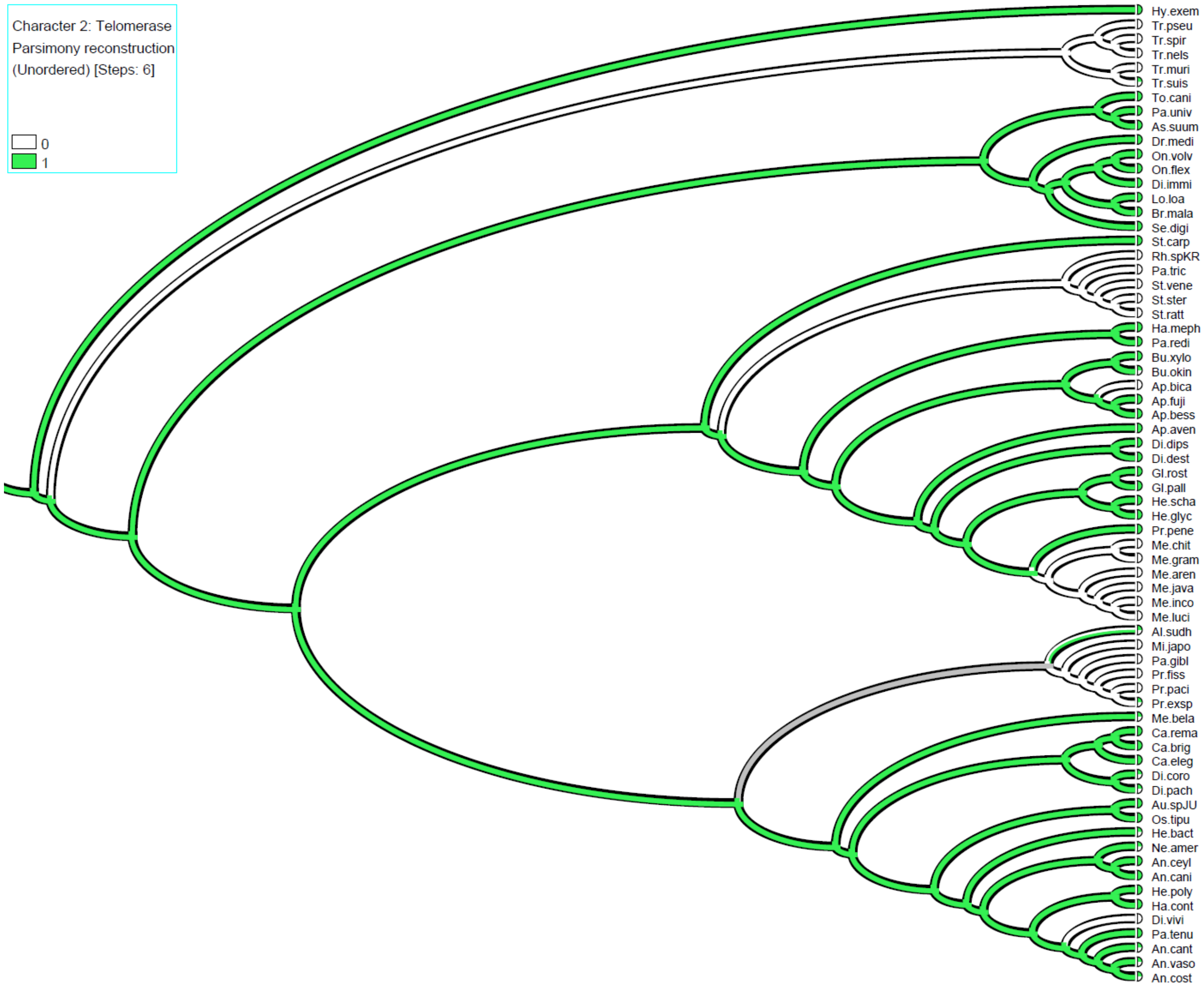

Based on presence / absence of the telomerase enzyme in nematode proteomes or telomerase gene in the genomes, ancestral states were reconstructed using parsimony in Mesquite (Maddison and Maddison, 2014). Green means telomerase present and white absent.

### Supplementary Figure 7: enriched motif at *M. incognita* contig ends

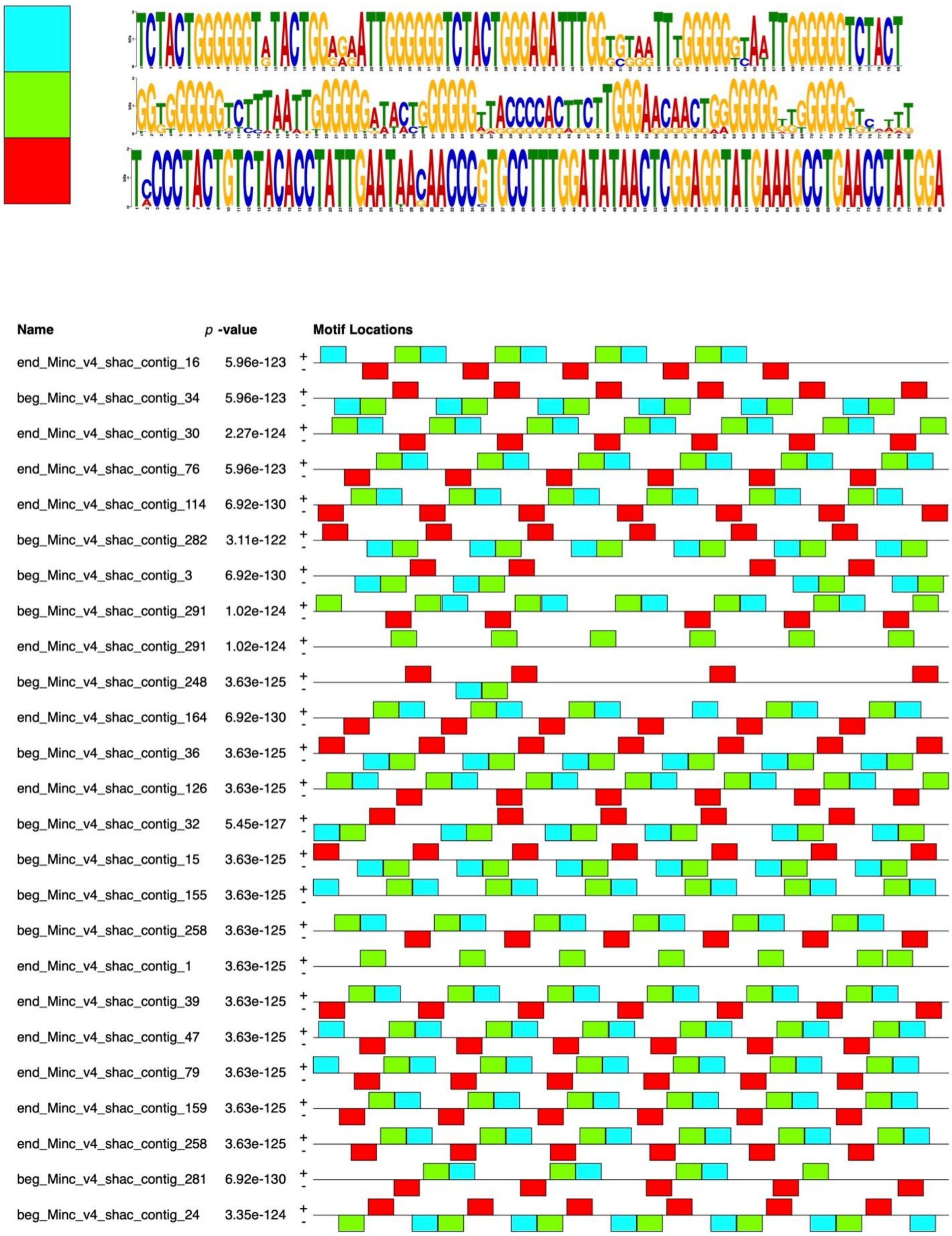

Enriched repeated motifs by increasing e-value and their distribution on contig extremities (first and last 2kb) in the *M. incognita* genome.

### Supplementary Figure 8: consensus of the enriched motif at *M. incognita* contig ends

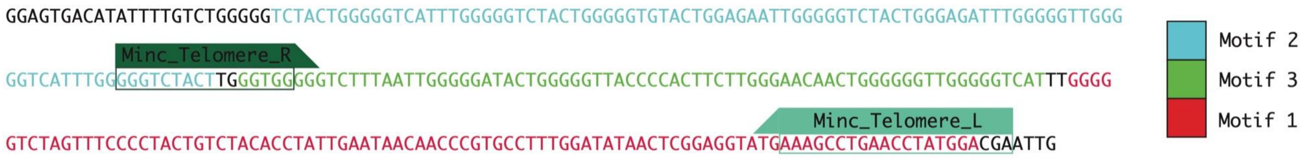

Positions of the primers used for PCR and FISH experiments are indicated as Minc_Telomere-R and Minc_Telomere_L

**GGAGTGACATATTTTGTCTGGGGGTCTACTGGGGGTCATTTGGGGGTCTACTGGGGGTGTACTGGAGAATTGGGGGTCTACTGGGAGATTTGGGGGTTGGGGGTCATTTGG*GGGTCTACTTGGGTGGG*GGTCTTTAATTGGGGGATACTGGGGGTTACCCCACTTCTTGGGAACAACTGGGGGGTTGGGGGTCATTTGGGGGTCTAGTTTCCCCTACTGTCTACACCTATTGAATAACAACCCGTGCCTTTGGATATAACTCGGAGGTATG*AAAGCCTGAACCTATGGACGA*ATTG**

*(in italics: positions of the primers)*

Multiple sequence alignment of the repeated unit made of motif-2 (blue), motif-3 (green), and motif-1 (red) as defined in supplementary Figure 7 allowed deducing a ca. 250 - 300 bp consensus sequence.

Supplementary Figure 9: distribution of repeat patterns and G4-quadruplex on *M. incognita* contigs

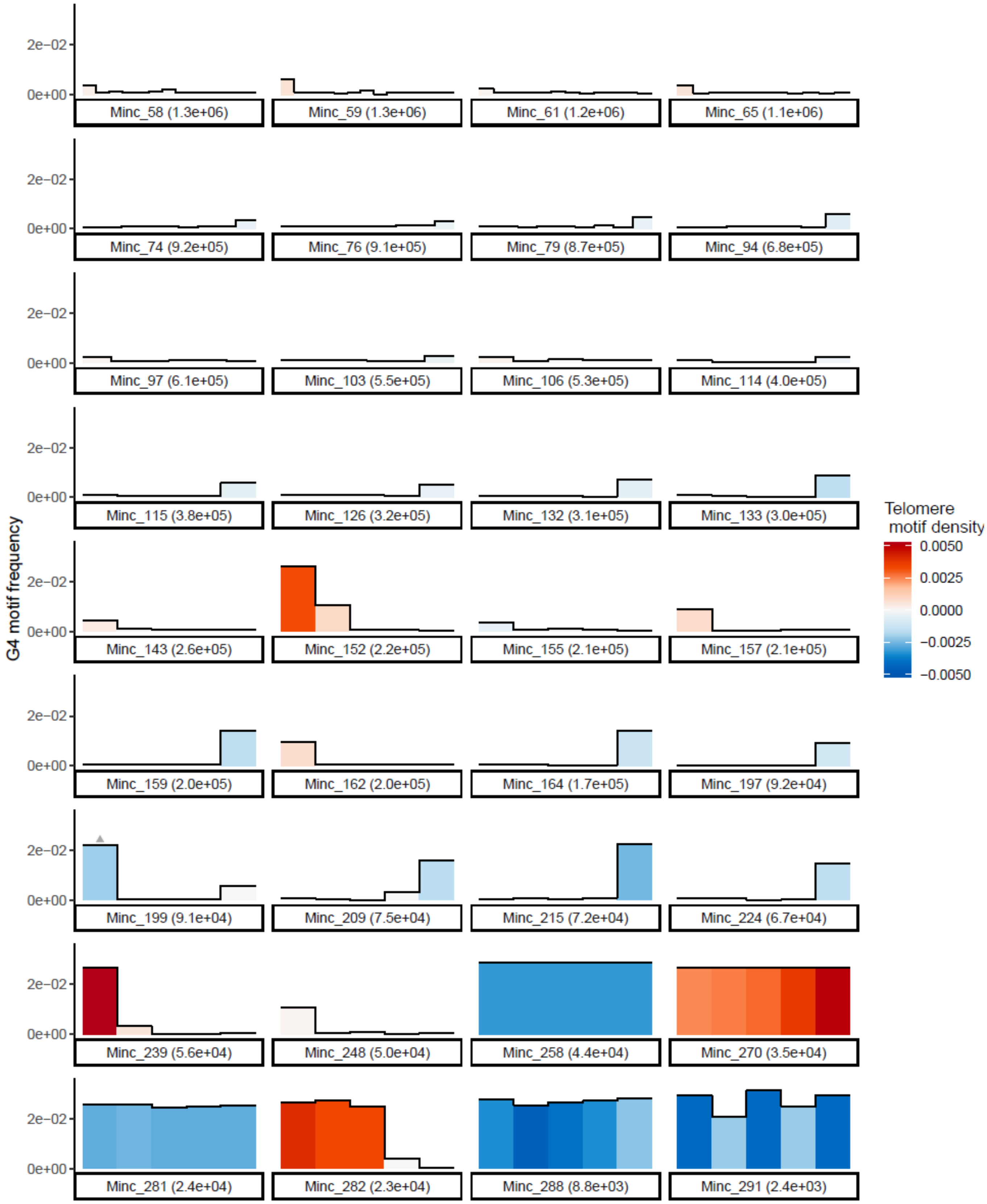

**Density of repeat patterns and G4-quadruplex along Minc contigs.** The density of repeated complex patterns is represented by a color gradient with positive values (red) indicating a density in the sense strand and negative values (blue) indicating a density on the reverse complement strand. Gray triangles above bars indicate regions of the contigs where less than 100% of the repeat patterns are on the same strand.The heights of bars in the histogram represent the density of G4-quadruplex forming regions in the same 100kb windows. Only contigs containing repeat arrays and smaller than the N80 value are represented, the same information for the bigger repeat-containing contigs is available in (Figure 3).

### Supplementary Table 3: position of Minc repeats on Minc contigs

Table available online at:
https://entrepot.recherche.data.gouv.fr/privateurl.xhtml?token=59e89b70-485f-4d0a-9e8d-cd1caa710037

### Supplementary Table 4: position of Minc repeats on Mare contigs

Table available online at:
https://entrepot.recherche.data.gouv.fr/privateurl.xhtml?token=d4224cdc-bab5-48bc-8507-ed0afddd6779

### Supplementary Table 5: position of Minc repeats on Mjav contigs

Table available online at:
https://entrepot.recherche.data.gouv.fr/privateurl.xhtml?token=84cd9e3a-f94b-4530-bfee-8536de453425

### Supplementary Figure 10: distribution of repeat patterns and G4-quadruplex on *M. javanica* contigs

**A )Consensus sequence of *M. javanica***

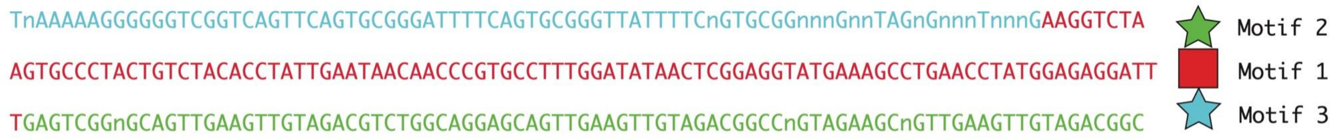

**TnAAAAAGGGGGGTCGGTCAGTTCAGTGCGGGATTTTCAGTGCGGGTTATTTTCnGTGCGGnnnGnnTAGn GnnnTnnnGAAGGTCTAAGTGCCCTACTGTCTACACCTATTGAATAACAACCCGTGCCTTTGGATATAACTC GGAGGTATGAAAGCCTGAACCTATGGAGAGGATTTGAGTCGGnGCAGTTGAAGTTGTAGACGTCTGGCAG GAGCAGTTGAAGTTGTAGACGGCCnGTAGAAGCnGTTGAAGTTGTAGACGGC**

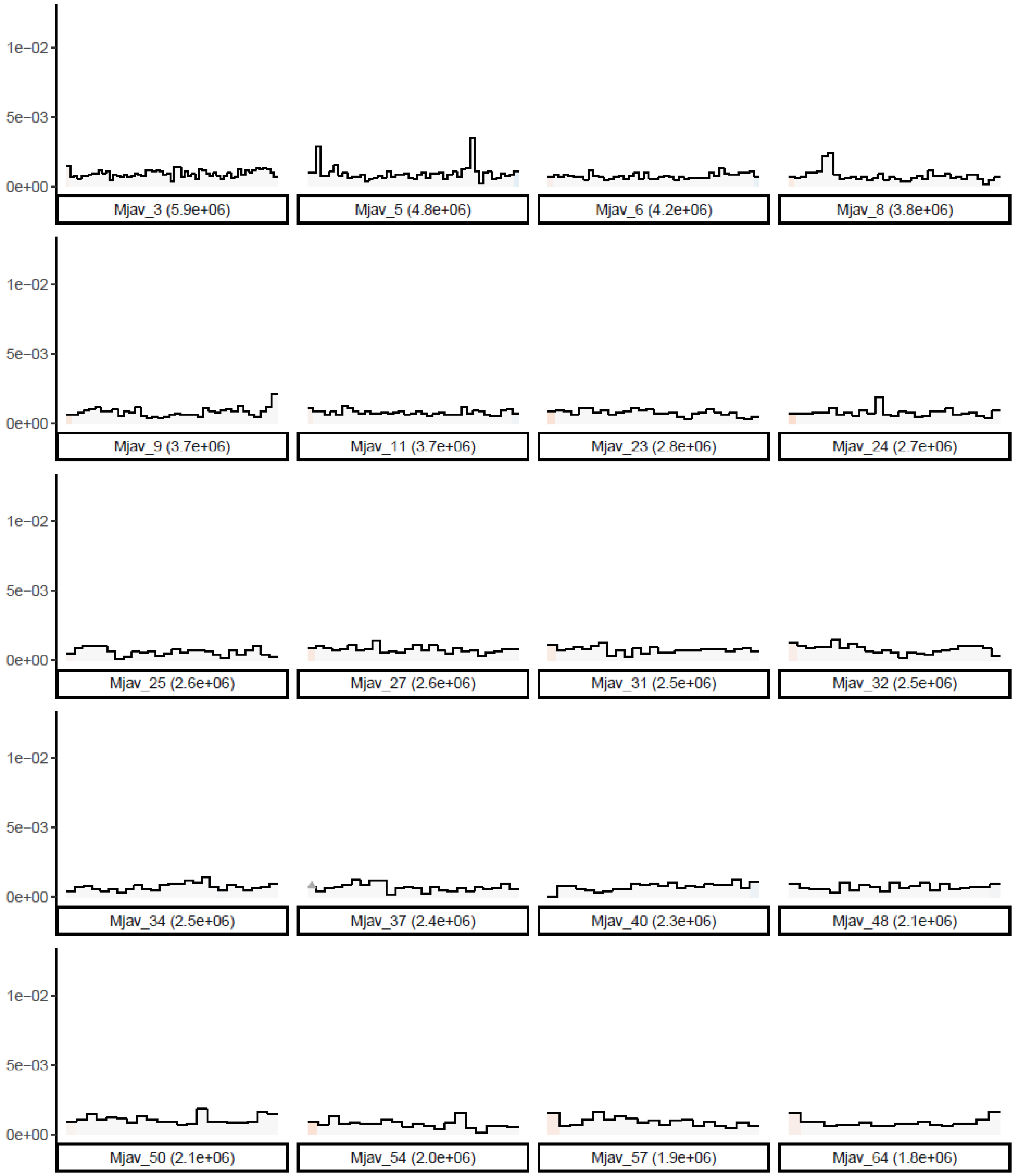
1e−02
5e−03
0e+00
Mjav_3 (5.9e+06)
Mjav_5 (4.8e+06)
Mjav_6 (4.2e+06)
Mjav_8 (3.8e+06)
Mjav_9 (3.7e+06)
Mjav_11 (3.7e+06)
Mjav_23 (2.8e+06)
Mjav_24 (2.7e+06)
Mjav_25 (2.6e+06)
Mjav_27 (2.6e+06)
Mjav_31 (2.5e+06)
Mjav_32 (2.5e+06)
Mjav_34 (2.5e+06)
Mjav_37 (2.4e+06)
Mjav_40 (2.3e+06)
Mjav_48 (2.1e+06)
Mjav_50 (2.1e+06)
Mjav_54 (2.0e+06)
Mjav_57 (1.9e+06)
Mjav_64 (1.8e+06)

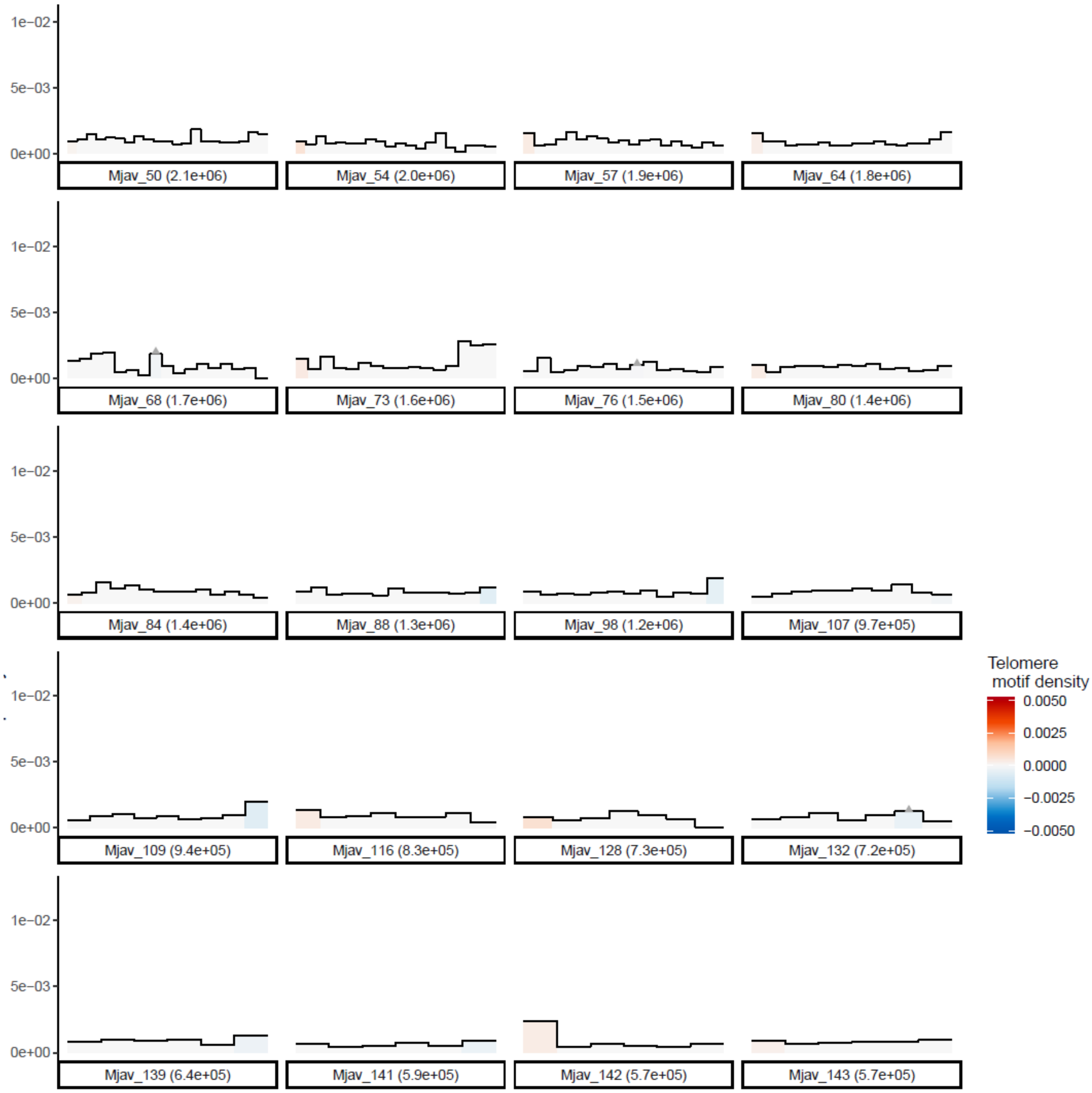
1e−02
5e−03
0e+00
Mjav_50 (2.1e+06)
Mjav_54 (2.0e+06)
Mjav_57 (1.9e+06)
Mjav_64 (1.8e+06)
Mjav_68 (1.7e+06)
Mjav_73 (1.6e+06)
Mjav_76 (1.5e+06)
Mjav_80 (1.4e+06)
Mjav_84 (1.4e+06)
Mjav_88 (1.3e+06)
Mjav_98 (1.2e+06)
Mjav_107 (9.7e+05)
Mjav_109 (9.4e+05)
Mjav_116 (8.3e+05)
Mjav_128 (7.3e+05)
Mjav_132 (7.2e+05)
Mjav_139 (6.4e+05)
Mjav_141 (5.9e+05)
Mjav_142 (5.7e+05)
Mjav_143 (5.7e+05)
Telomere motif density
0.0050
0.0025
0.0000
−0.0025
−0.0050

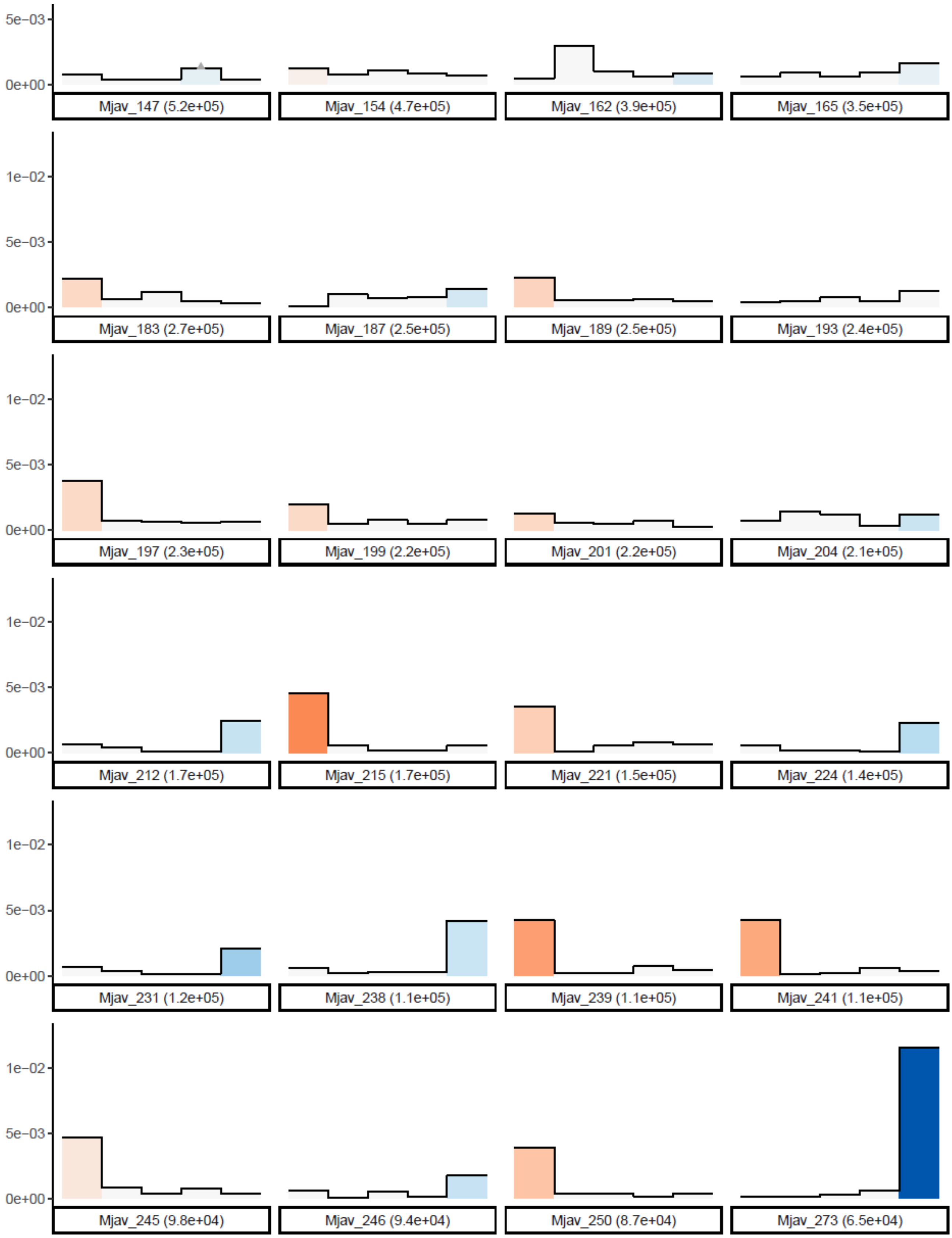

**B) Density of repeat patterns and G4-quadruplex along *M. javanica* contigs.** The density of repeated complex patterns is represented by a color gradient with positive values (red) indicating a density in the sense strand and negative values (blue) indicating a density on the reverse complement strand. Gray triangles above bars indicate regions of the contigs where less than 100% of the repeat patterns are on the same strand.The heights of bars in the histogram represent the density of G4-quadruplex forming regions in the same 100kb windows.

### Supplementary Figure 11: distribution of repeat patterns and G4-quadruplex on *M. arenaria* contigs

#### A) Consensus sequence of *M. arenaria*

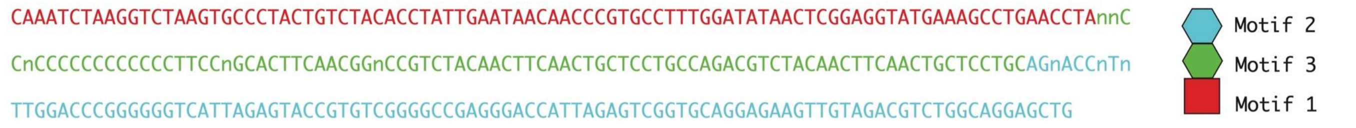

**CAAATCTAAGGTCTAAGTGCCCTACTGTCTACACCTATTGAATAACAACCCGTGCCTTTGGATATAACTCGG AGGTATGAAAGCCTGAACCTAnnCCnCCCCCCCCCCCCTTCCnGCACTTCAACGGnCCGTCTACAACTTC AACTGCTCCTGCCAGACGTCTACAACTTCAACTGCTCCTGCAGnACCnTnTTGGACCCGGGGGGTCATTAG AGTACCGTGTCGGGGCCGAGGGACCATTAGAGTCGGTGCAGGAGAAGTTGTAGACGTCTGGCAGGAGCT G**

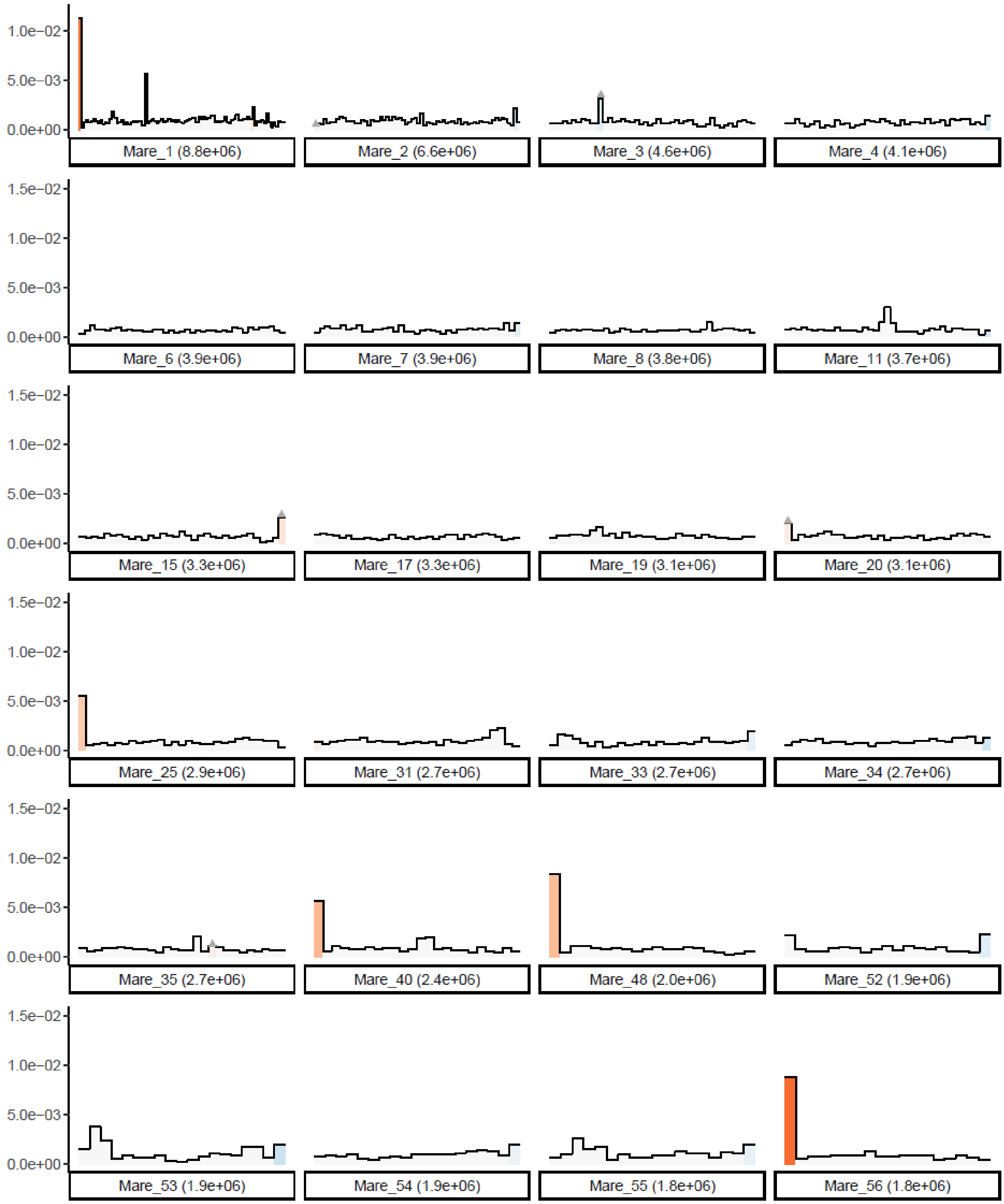
1.0e−02
5.0e−03
0.0e+00
1.5e−02
Mare_1 (8.8e+06)
Mare_2 (6.6e+06)
Mare_3 (4.6e+06)
Mare_4 (4.1e+06)
Mare_6 (3.9e+06)
Mare_7 (3.9e+06)
Mare_8 (3.8e+06)
Mare_11 (3.7e+06)
Mare_15 (3.3e+06)
Mare_17 (3.3e+06)
Mare_19 (3.1e+06)
Mare_20 (3.1e+06)
Mare_25 (2.9e+06)
Mare_31 (2.7e+06)
Mare_33 (2.7e+06)
Mare_34 (2.7e+06)
Mare_35 (2.7e+06)
Mare_40 (2.4e+06)
Mare_48 (2.0e+06)
Mare_52 (1.9e+06)
Mare_53 (1.9e+06)
Mare_54 (1.9e+06)
Mare_55 (1.8e+06)
Mare_56 (1.8e+06)

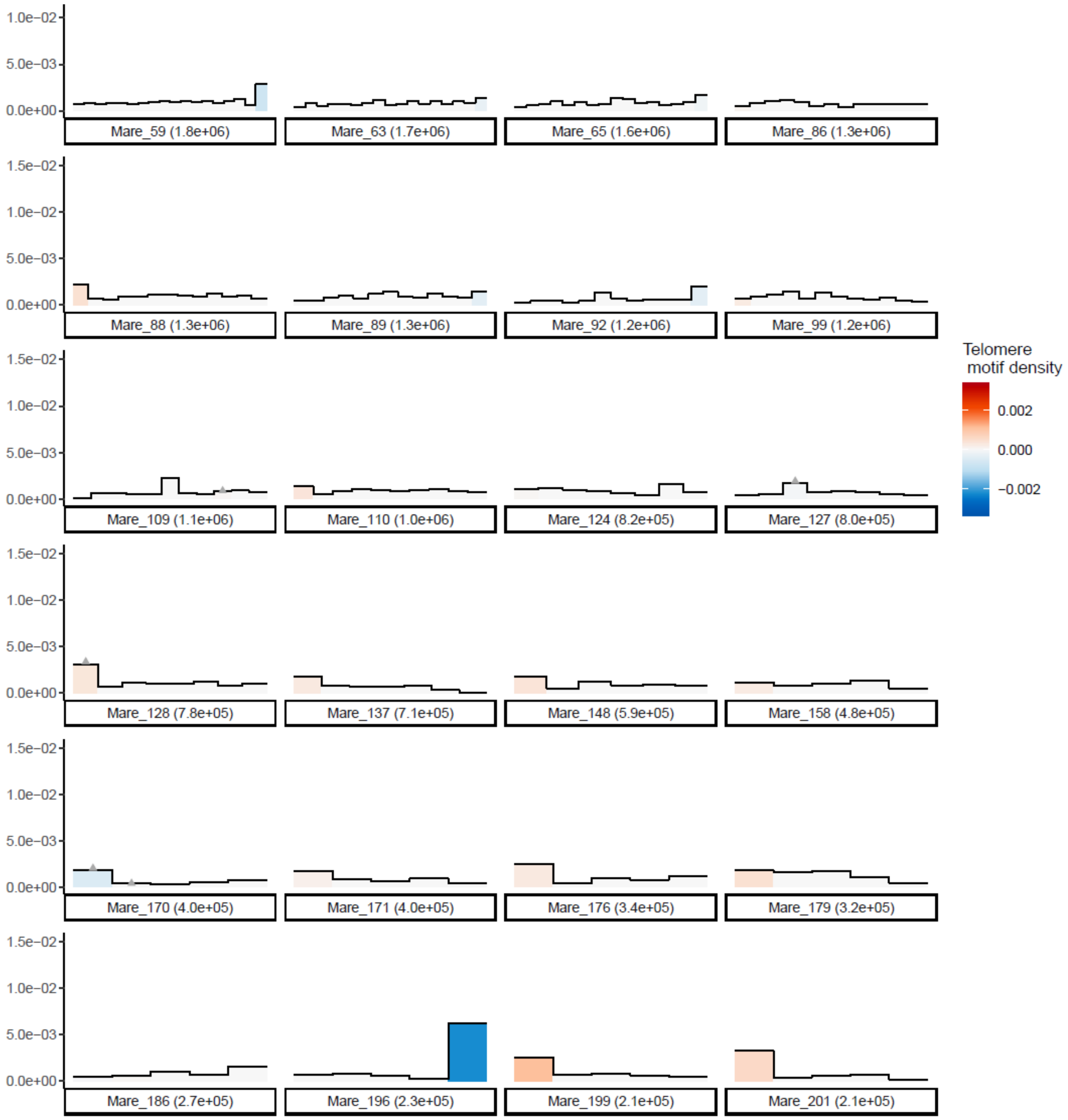
Mare_59 (1.8e+06)
Mare_63 (1.7e+06)
Mare_65 (1.6e+06)
Mare_86 (1.3e+06)
Mare_88 (1.3e+06)
Mare_89 (1.3e+06)
Mare_92 (1.2e+06)
Mare_99 (1.2e+06)
Mare_109 (1.1e+06)
Mare_110 (1.0e+06)
Mare_124 (8.2e+05)
Mare_127 (8.0e+05)
Mare_128 (7.8e+05)
Mare_137 (7.1e+05)
Mare_148 (5.9e+05)
Mare_158 (4.8e+05)
Mare_170 (4.0e+05)
Mare_171 (4.0e+05)
Mare_176 (3.4e+05)
Mare_179 (3.2e+05)
Mare_186 (2.7e+05)
Mare_196 (2.3e+05)
Mare_199 (2.1e+05)
Mare_201 (2.1e+05)
Telomere motif density
0.002
0.000
−0.002
1.5e−02
1.0e−02
5.0e−03
0.0e+00

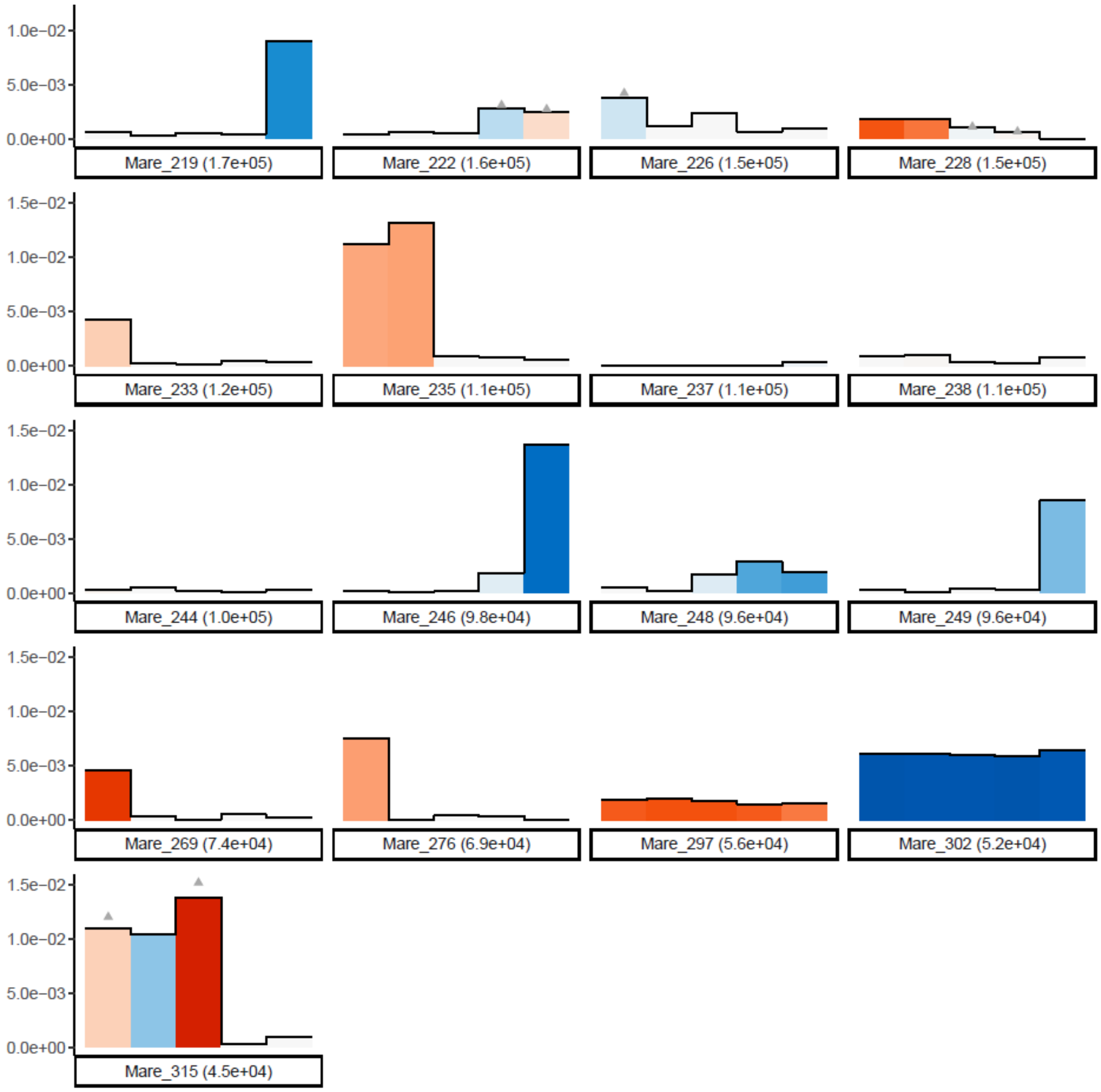

**B) Density of repeat patterns and G4-quadruplex along *M. arenaria* contigs.** The density of repeated complex patterns is represented by a color gradient with positive values (red) indicating a density in the sense strand and negative values (blue) indicating a density on the reverse complement strand. Gray triangles above bars indicate regions of the contigs where less than 100% of the repeat patterns are on the same strand.The heights of bars in the histogram represent the density of G4-quadruplex forming regions in the same 100kb windows.

Frequency of G4-quadruplexes along contigs with predicted telomeric sequences in *M. arenaria*.

### Supplementary Table 6: position of Minc repeats on Mluci contigs

Table available online at:

https://entrepot.recherche.data.gouv.fr/privateurl.xhtml?token=6d137b90-3599-4134-a75d-3a0679be7448

### Supplementary Table 7: annotation of (retro)transposons and other repeats

The annotation of transposable elements and other repetitive regions on the genomes of *M. incognita*, *M. arenaria* and *M. javanica* was performed using EDTA (Ou et al., 2019) version 2.1.0.

| | | ***M. incognita*** | | ***M. arenaria*** | | ***M. javanica*** | |
|---|---|---|---|---|---|---|---|
| Class | Type | Count | bpMasked | Count | bpMasked | Count | bpMasked |
| **Retrotransposons** | | | | | | | |
| LTR | Copia | 25 | 8,783 | 444 | 185,968 | 290 | 134,146 |
| | Gypsy | 10216 | 5,548,981 | 12585 | 12,152,090 | 9704 | 8,940,100 |
| | Unknown | 11048 | 6,556,744 | 25891 | 11,396,311 | 11646 | 5,609,431 |
| nonLTR | LINE_element | 370 | 373,486 | 503 | 793,499 | 528 | 703,477 |
| **DNA Transposons** | | | | | | | |
| TIR | CACTA | 4296 | 896,156 | 5464 | 1,186,379 | 4370 | 1,113,443 |
| | Mutator | 30418 | 7,227,848 | 48371 | 12,631,831 | 54636 | 17,274,061 |
| | PIF_Harbinger | 2231 | 522,128 | 2513 | 696,,702 | 2096 | 531,815 |
| | Tc1_Mariner | 476 | 178,149 | 2128 | 522,,780 | 1963 | 524,585 |
| | hAT | 55062 | 11,913,136 | 94287 | 22,654,778 | 85261 | 19,036,926 |
| | polinton | 2561 | 2,988,992 | 1755 | 1,709,300 | 4784 | 5,536,271 |
| nonTIR | helitron | 51164 | 2,635,069 | 4369 | 4,726,375 | 4077 | 5,409,339 |
| **Unclassified / unknown repeats** | | | | | | | |
| repeat region | repeat | 27,789 | 8,705,800 | 45567 | 19,646,238 | 55511 | 19,620,486 |

### Supplementary Figure 12. Gel electrophoresis of amplified telomere sequence.

A) Gradient PCR using primer pair (Supplementary Figure 12) and temperature range as listed in Materials and Methods section. B) Labeled probe for telomere sequence using biotin-dUTP nucleotide in PCR reaction.

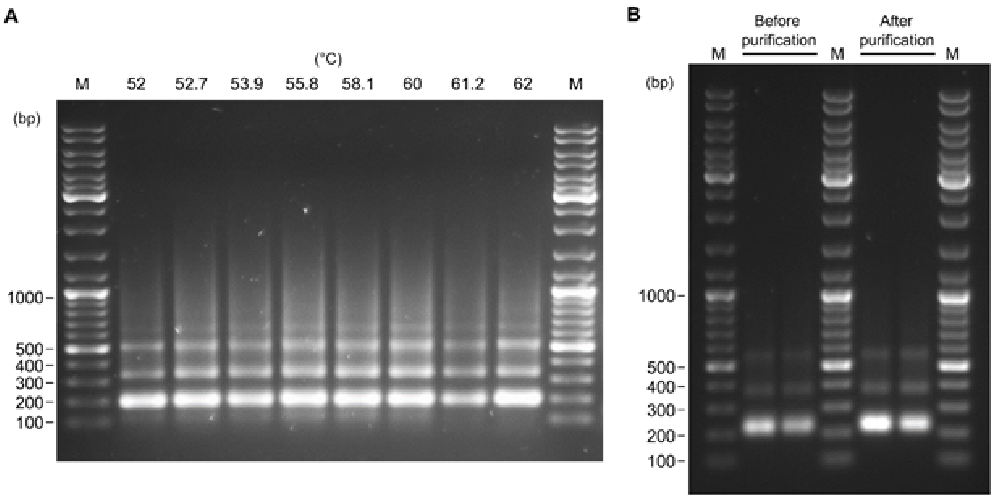

### Supplementary Table 8: de novo assembled Minc transcripts containing the repeat and matching the genome

Table available online at:
https://entrepot.recherche.data.gouv.fr/privateurl.xhtml?token=8cb3f59d-ffe8-4af4-8079-1db1cc4bc9b2

### Supplementary Table 9: ISO-seq Minc transcripts containing the repeat and matching the genome

Table available online at:

https://entrepot.recherche.data.gouv.fr/privateurl.xhtml?token=dd74a782-df20-418e-8462-7eddf8b67a51

### Supplementary Table 10: Raw ONT sequencing library information.

Sequencing libraries were produced using the Minion or Promethion platform, with flow-cell version 9.4.1 in both cases. Libraries sequenced with a Minion device were then base-called using guppy dna_r9.4.1_450bps_sup.cfg configuration file. Libraries sequenced with a Promethion device were then base-called using guppy dna_r9.4.1_450bps_sup_prom.cfg configuration file. All base calling was made in super high accuracy mode.

| Species | Raw sequencing library | Platform | Accession numbers |
|---|---|---|---|
| *M. incognita* | BZU_AD_ONT_1_FAK08876_A | Minion | ERX10638972 |
| *M. incognita* | BZU_AA_ONT_1_FAH89151_A | Minion | ERX10638973 |
| *M. incognita* | BZU_AA_ONT_1_FAF18555_A | Minion | ERX10638974 |
| *M. incognita* | BZU_AA_ONT_1_FAH58676_A | Minion | ERX10638975 |
| *M. incognita* | BZU_AA_ONT_1_FAH54346_A | Minion | ERX10638976 |
| *M. incognita* | BZU_AA_ONT_1_FAH43582_A | Minion | ERX10638977 |
| *M. incognita* | BZU_AA_ONT_1_FAH54287_A | Minion | ERX10638978 |
| *M. incognita* | BZU_AA_ONT_1_FAH63986_A | Minion | ERX10638979 |
| *M. incognita* | BZU_AA_ONT_1_FAH60611_A | Minion | ERX10638980 |
| *M. incognita* | BZU_AA_ONT_1_PAD23691_A | Promethion | ERX10638981 |
| *M. incognita* | BZU_AA_ONT_1_PAC22366_A | Promethion | ERX10638982 |
| *M. javanica* | BZU_AH_ONT_1_FAK26065_A | Minion | ERX10638983 |
| *M. javanica* | BZU_AC_ONT_1_FAH59483_A | Minion | ERX10638984 |
| *M. javanica* | BZU_AC_ONT_1_FAH54228_A | Minion | ERX10638985 |

| | | | |
|---|---|---|---|
| *M. javanica* | BZU_AC_ONT_1_FAH43433_A | Minion | ERX10638986 |
| *M. javanica* | BZU_AC_ONT_1_FAH58655_A | Minion | ERX10638987 |
| *M. javanica* | BZU_AC_ONT_1_FAH54419_A | Minion | ERX10638988 |
| *M. javanica* | BZU_AG_ONT_1_PAD26893_A | Promethion | ERX10638989 |
| *M. javanica* | BZU_AC_ONT_1_PAC15472_A | Promethion | ERX10638990 |
| *M. arenaria* | BZU_AH_ONT_1_PAD26768_A | PromethION | ERX10639152 |
| *M. arenaria* | BZU_AG_ONT_1_FAK26121_A | Minion | ERX10638991 |
| *M. arenaria* | BZU_AB_ONT_1_FAH54179_A | Minion | ERX10638992 |
| *M. arenaria* | BZU_AB_ONT_1_FAH58521_A | Minion | ERX10638993 |
| *M. arenaria* | BZU_AB_ONT_1_FAH58614_A | Minion | ERX10638994 |
| *M. arenaria* | BZU_AB_ONT_1_FAH54190_A | Minion | ERX10638995 |
| *M. arenaria* | BZU_AB_ONT_1_FAH43612_A | Minion | ERX10638996 |
| *M. arenaria* | BZU_AB_ONT_1_FAH59474_A | Minion | ERX10638997 |

### Supplementary Table 11: Raw Illumina data used for genome polishing and k-mer analyses

| Species | Raw sequencing library | Reads | Length | Accession numbers |
|---|---|---|---|---|
| *M. incognita* | BZU_AAOSDE_1_1_H732LBCX2.12BA290_clean.fastq | 19,805,212 | 250 | ERX10638998 |
| *M. incognita* | BZU_AAOSDE_1_2_H732LBCX2.12BA290_clean.fastq | 19,805,212 | 250 | ERX10638998 |
| *M. incognita* | BZU_AAOSDE_2_1_H732LBCX2.12BA290_clean.fastq | 20,090,514 | 250 | ERX10638999 |
| *M. incognita* | BZU_AAOSDE_2_2_H732LBCX2.12BA290_clean.fastq | 20,090,514 | 250 | ERX10638999 |
| *M. javanica* | BZU_ACOSDE_2_1_H52YLBCX2.12BA292_clean.fastq | 64,456,448 | 250 | ERX10639000 |
| *M. javanica* | BZU_ACOSDE_2_2_H52YLBCX2.12BA292_clean.fastq | 64,456,448 | 250 | ERX10639000 |
| *M. arenearia* | BZU_ABOSDE_1_1_H732LBCX2.12BA291_clean.fastq | 27,049,894 | 250 | ERX10639001 |
| *M. arenearia* | BZU_ABOSDE_1_2_H732LBCX2.12BA291_clean.fastq | 27,049,894 | 250 | ERX10639001 |
| *M. arenearia* | BZU_ABOSDE_2_1_H732LBCX2.12BA291_clean.fastq | 27,421,721 | 250 | ERX10639002 |
| *M. arenearia* | BZU_ABOSDE_2_2_H732LBCX2.12BA291_clean.fastq | 27,421,721 | 250 | ERX10639002 |
